# Deep Visual Proteomics links vascular smooth muscle cell phenotypes to atherosclerotic plaque stability

**DOI:** 10.64898/2026.07.17.738909

**Authors:** Elena Kratz, Ankit Sinha, Trusha Adeshara, Valentina Paloschi, Nadja Sachs, Jessica Pauli, Nadyia Glukha, Annalena Huber, Sara Schanda, Andreas Metousis, Tim Heymann, Daniela Branzan, Tore Bleckwehl, Sikander Hayat, Lars Maegdefessel, Matthias Mann

**Author notes:** Shared corresponding authors.

## Abstract

Vascular smooth muscle cells (VSMCs) drive atherosclerosis through phenotypic switching, yet their spatial organization and protein signatures within plaques remain poorly characterized. Here, we applied Deep Visual Proteomics (DVP) to dissect more than 500 VSMC neighborhoods across 24 human carotid plaques and profile VSMC plasticity in disease. To functionally interpret these tissue proteomes, we built a reference atlas of primary VSMCs driven toward five phenotypes by TGF-β, PDGF-BB, osteogenic stimuli, IL-1β, or cholesterol, quantifying over 10,000 proteins. We integrated tissue and reference proteomes with a deep-learning framework that assigns functional phenotypes to each neighborhood. This revealed spatially distinct phenotype distributions and a shift toward dedifferentiated states in unstable plaques. Knockdown of four candidates (TNC, TNFAIP2, AEBP1, PLK1) validated operational roles during phenotypic switching. Our approach functionally annotates spatial proteomes and links VSMC plasticity to plaque instability in carotid artery disease.

## Introduction

Atherosclerosis is the main driver of cardiovascular diseases, the leading cause of death worldwide^1^. It is a chronic, lipid-driven inflammatory disease characterized by progressive plaque formation that leads to vessel stenosis, and to acute ischemic complications when plaques become unstable and rupture^2^. Atherosclerotic plaques are highly heterogeneous structures composed of vascular smooth muscle cells (VSMCs), endothelial cells and immune cells, embedded within an extracellular matrix (ECM), alongside calcified deposits and lipid cores^3,4^. As the predominant cell type of the healthy arterial wall and comprising up to 70% of plaque cells, VSMCs have emerged as key regulators of plaque initiation, progression and stability^5–7^. In healthy vessels, VSMCs adopt a quiescent, contractile phenotype that maintains vascular tone and structural integrity. In response to atherogenic stimuli, such as inflammatory signals, oxidized lipids and disturbed flow, VSMCs undergo phenotypic switching to myofibroblast-like, proliferative, osteogenic and foam cell-like states^8–10^. This plasticity enables them to drive ECM remodeling, calcification, inflammatory signaling and plaque expansion, thereby shaping plaque architecture and clinical outcomes^11,12^.

Despite the recognized importance of VSMCs in plaque dynamics, their spatial organization and molecular phenotypes within atherosclerotic plaques remain poorly characterized. Previous efforts to understand VSMC phenotypic diversity in human plaques mainly relied on single-cell RNA sequencing, which uncovered multiple dedifferentiated states in atherosclerotic plaques^13,14^. More recently, spatial transcriptomics added cellular resolution *in situ*, preserving the tissue architecture^15,16^. However, RNA-based approaches cannot capture protein-level programs that ultimately drive VSMC function. Proteomics provides a direct readout of these programs, and we recently applied histomorphology-guided spatial proteomics to resolve subregion-specific signatures of plaque instability in human carotid plaques^17^. Subregion-level analysis, however, aggregates heterogeneous cell populations and cannot resolve individual cellular phenotypes. Deep Visual Proteomics (DVP) is an emerging approach that resolves the proteome of various cell types within formalin-fixed paraffin-embedded (FFPE) tissues^18^. By combining immunofluorescence (IF) imaging, machine learning-based cell segmentation, and laser microdissection of target cells with ultra-sensitive mass spectrometry, DVP enables comprehensive proteomic profiling of defined cell populations within intact histological sections while preserving the spatial context.

In this study, we applied DVP to map spatial proteomes of VSMC-enriched neighborhoods in human carotid plaques, thereby capturing the proteomic landscape that defines VSMC diversity. To assign functional traits to these spatial proteomes, we developed a deep learning framework that combines DVP-derived proteomes with reference proteomes from primary VSMC phenotype models. This strategy allowed us to identify pathway alterations underlying VSMC phenotypic transitions and uncovered spatially restricted proteomic signatures that correlate with plaque instability.

## Results

### DVP captures VSMC proteomes from human atherosclerotic plaques

Using DVP, we analyzed 24 FFPE human atherosclerotic carotid plaques. The cohort comprised equal numbers of stable (n = 12) and unstable (n = 12) plaques, with clinical covariates balanced across groups (Extended Figure 1a, Extended Data Table 1). Plaque stability status was defined by fibrous cap thickness, with unstable defined as < 200 µm at the thinnest region in accordance with the criteria set by the Oxford Plaque Studies for carotid artery disease^19^. An additional characterization of plaque status was performed following the guidelines established by the American Heart Association (AHA)^20^. To ensure adequate spatial sampling, we selected plaques with intact tunica media and abundant cellular content based on hematoxylin/eosin (HE) and Elastica van Gieson (EvG) staining.

To benchmark the DVP workflow on atherosclerotic plaque tissue, we laser-microdissected three different cell types (VSMCs, macrophages, T cells) from two representative carotid plaques (ID7, ID21), revealing clear separation of the three populations and selective enrichment of canonical marker proteins (Extended Figures 1b-d). We then implemented a tailored DVP pipeline to profile VSMC neighborhoods from plaques (Figure 1a). VSMCs were stained by immunofluorescence using a smooth muscle actin (ACTA2) antibody, followed by segmentation of ACTA2+ cells. Then, we applied a two-step clustering algorithm, combining supervised centroid placement with density-based clustering (HDBSCAN), to group ACTA2+ cells into spatial neighborhoods. To account for inter-plaque variability of ACTA2+ cell area and location, we supervised our approach by manually defining count and location of neighborhood centroids (Extended Figure 1e). For each neighborhood, we laser-microdissected 400 ACTA2+ shapes, yielding 545 neighborhood samples from 24 plaques, which were analyzed by Data-Independent Acquisition (DIA) on an Orbitrap Astral mass spectrometer at 80 samples per day.

**Figure 1:**
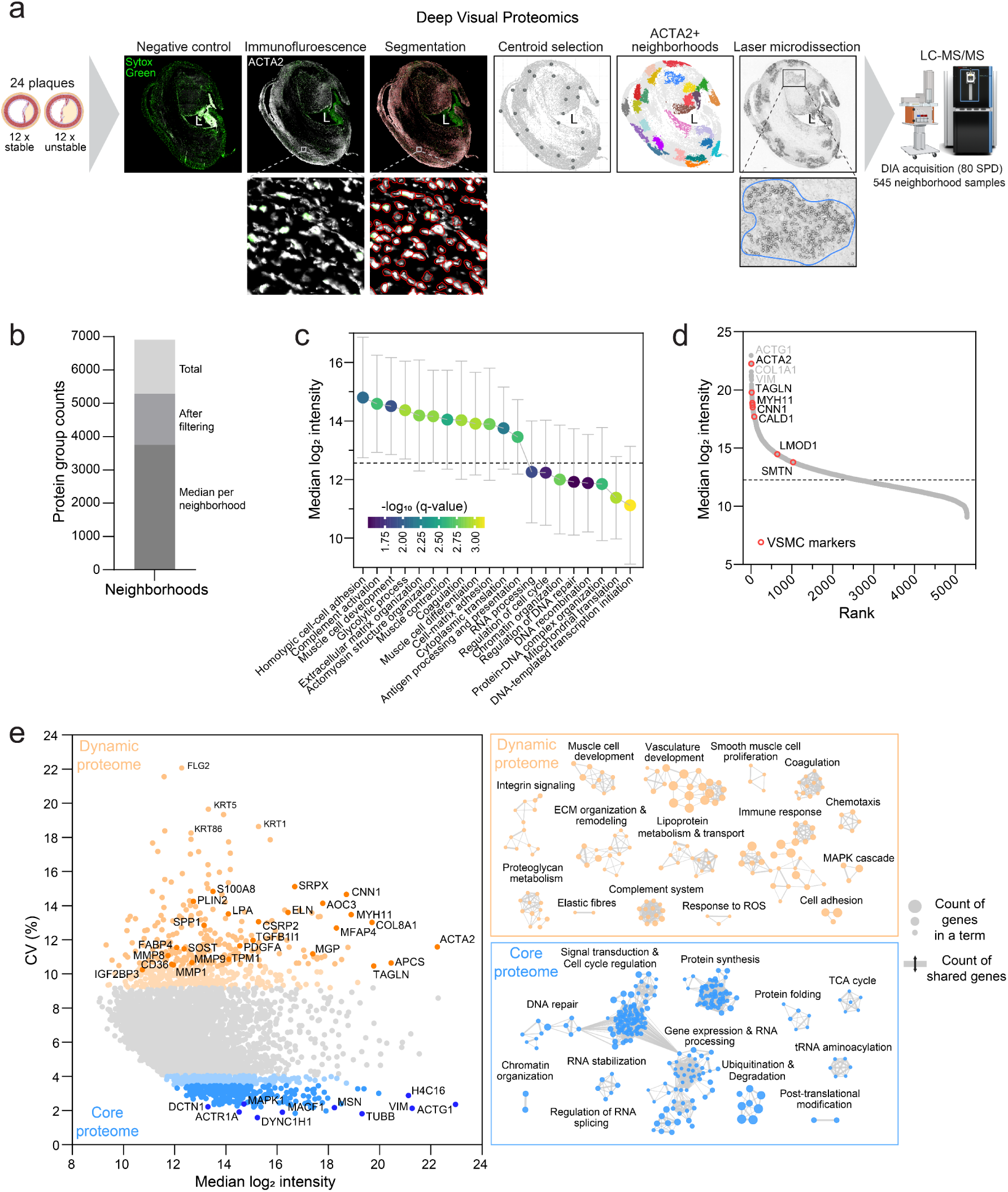
DVP captures VSMC neighborhood proteomes from human atherosclerotic plaques. **a** DVP workflow applied to 24 FFPE human carotid plaques. Sections were stained with an anti-ACTA2 antibody and Sytox Green, ACTA2+ shapes were segmented and clustered into neighborhoods of 400 ACTA2+ shapes each (approx. 28,000 µm^2^ per neighborhood), 545 neighborhoods were laser-microdissected, and acquired by liquid chromatography-mass spectrometry (LC-MS/MS) in Data-Independent Acquisition (DIA) mode at 80 samples per day (SPD), L = plaque lumen. **b** Protein group counts across neighborhoods after applying sample-level quality filtering (535 of 545 neighborhoods retained). **c** Gene Ontology terms enriched in the filtered proteome (5,287 protein groups), ranked by median log_2_ intensity. The dashed line marks the median log_2_ intensity across all samples and proteins. **d** Rank-abundance plot of all filtered protein groups. Hallmark VSMC markers are highlighted in red. The dashed line marks the median log_2_ intensity across all samples and proteins. **e** Biological coefficient of variation (CV) versus median log_2_ intensity for each protein group. The 500 most variable proteins (dynamic proteome, orange) and 500 least variable proteins (core proteome, blue) are highlighted. Enriched biological pathways for each set are displayed as network plots.

After applying sample-level quality filtering, 535 of 545 neighborhoods were retained, yielding a total of 6,910 protein groups with a median of 3,754 proteins per neighborhood. After protein-level filtering, 5,287 protein groups were retained for downstream analysis (Figure 1b).

Subcellular compartment analysis across the whole dataset showed the expected proteomic architecture of VSMC-enriched plaque samples, dominated by cytoplasmic, endoplasmic reticulum (ER), extracellular and blood-derived proteins (Extended Figure 1f). Gene set enrichment analysis of overall protein abundance confirmed that many of the top-enriched terms reflected core VSMC biology, including muscle cell development, contraction and differentiation, alongside general cellular processes (cell-cell adhesion, glycolysis, translation) (Figure 1c). Consistent with the close association of VSMCs with their surrounding matrix, ECM-related processes were also among the enriched terms. Vascular– and blood-associated processes ranked highly as well, reflecting the direct exposure of plaque tissue to circulating blood, especially in advanced and unstable lesions. In contrast, transcription– and nucleus-associated terms were detected at notably lower abundance, consistent with their general underrepresentation in bottom-up proteomics workflows^21^.

The proteome data spanned a dynamic range of 4.5 orders of magnitude (Figure 1d). Hallmark VSMC markers, such as ACTA2, transgelin (TAGLN) and myosin heavy chain 11 (MYH11) were ranked in the top 10% of protein intensities across all neighborhoods and were consistently quantified in all neighborhood samples (Extended Figure 1g). These findings support enrichment of VSMC-specific proteomes and the specificity of our targeted cell type isolation.

To assess presence of phenotypic switching in our data, we quantified protein-level variation across all VSMC neighborhoods using the biological coefficient of variation (CV) (Figure 1e). Contractile proteins (ACTA2, TAGLN, MYH11) had large variation (top 10% of most variable proteins), along with markers of dedifferentiation, including ECM-interacting proteins (SRPX, COL8A1, ELN), matrix metalloproteinases (MMP1, MMP8, MMP9), lipid metabolism-associated proteins (PLIN2, FABP4, LPA), calcification-related proteins (SPP1, MGP, SOST) and markers of inflammation (AOC3, APCS, S100A8). We further examined the functional foundations of these variance patterns using gene set enrichment analysis (GSEA). Proteins with high variation (dynamic proteome) were enriched for muscle cell development, proliferation, ECM remodeling and immune-related processes. In contrast, proteins with low variation (core proteome) were dominated by housekeeping functionalities, such as cell cycle regulation, DNA repair and protein folding. The clear distinction between dynamic and core proteome provides molecular evidence for phenotypic plasticity within dissected VSMC neighborhoods of advanced human atherosclerotic plaques.

### DVP reveals molecular heterogeneity of VSMCs

To elucidate the molecular heterogeneity of VSMC neighborhoods, we performed principal component analysis (PCA) across all 535 neighborhoods (Figure 2a). Neighborhood samples arranged along a continuous trajectory in principal component space. ACTA2, a key contractile marker downregulated during phenotypic switching, showed a strong gradient along PC1, which was confirmed by the corresponding loadings (Figure 2b). Other contractile proteins (CNN1, MYH11, TAGLN) showed the same negative correlation with PC1, while dedifferentiation markers (e.g., LGALS3, MMP9, ABCA1, SPP1) were positively correlated with PC1. Enrichment analysis of high PC1-loading proteins identified lipid metabolism, translation, antigen presentation and stress response as the dominant programs (Extended Figure 2b). Together, this establishes a contractile-to-dedifferentiated continuum as the principal axis of heterogeneity within our neighborhood samples. To exclude technical confounders, we verified that this axis was not driven by patient identity (Extended Figure 2a) or blood contamination (Extended Figures 2c-e). Blood-derived proteins loaded primarily on PC2 and PC3, and blood-only PC1 values showed minor correlation with global PC1 values (Spearman ρ = 0.194, p-value < 0.001), suggesting that blood signal contributes an orthogonal dimension rather than driving the phenotypic ACTA2 axis.

**Figure 2:**
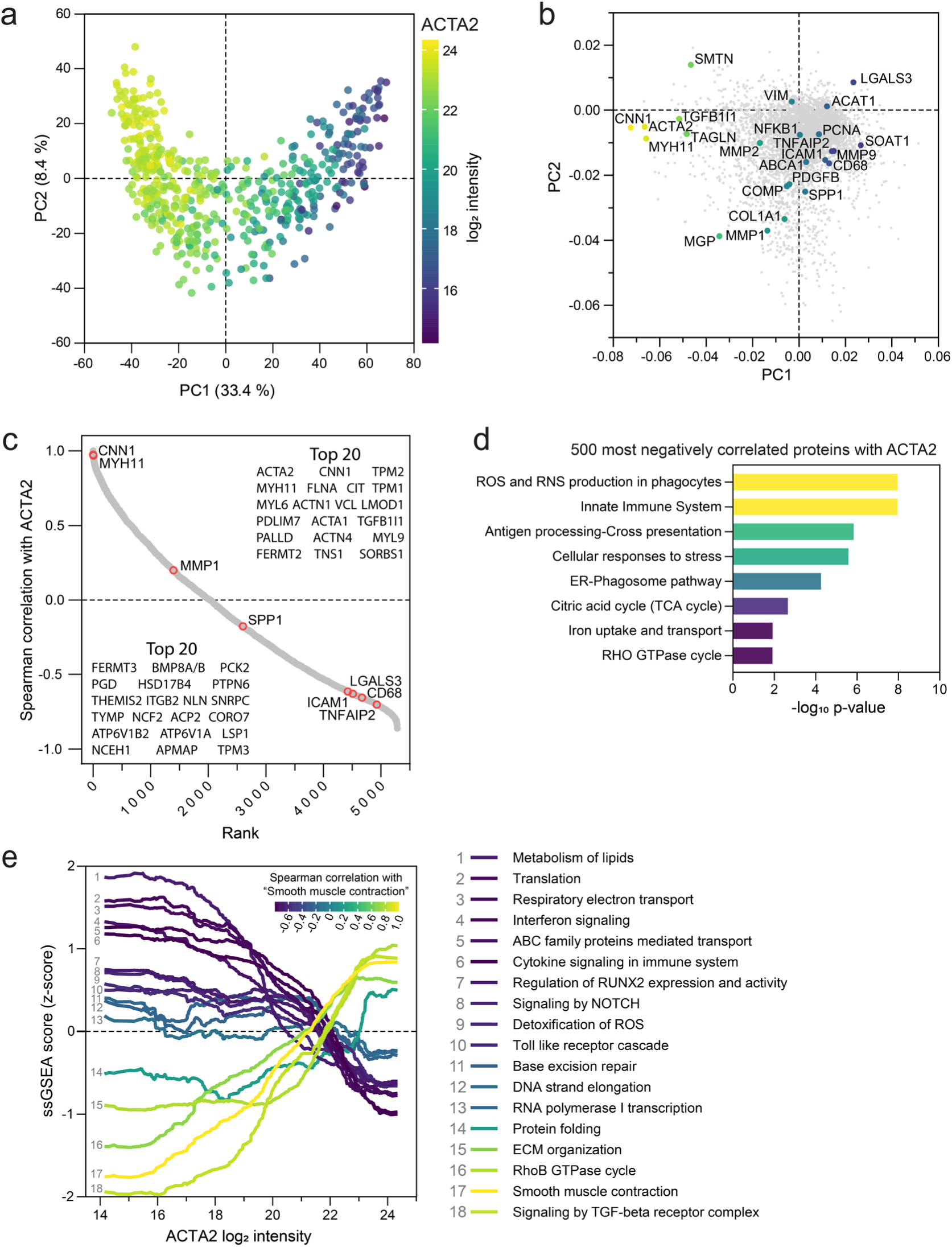
DVP reveals molecular heterogeneity of VSMCs. **a** PCA of VSMC neighborhood proteomes (n = 535), colored by ACTA2 log_2_ intensity. **b** PC1/PC2 loadings, selected markers of contractile and dedifferentiated VSMC states are labeled and colored by PC1 value. **c** Rank plot of the Spearman correlation between each protein’s log_2_ intensity and ACTA2. The top 20 most positively (top) and most negatively (bottom) correlated proteins are listed. **d** Over-representation analysis of the 500 proteins most negatively correlated with ACTA2. Selected Reactome terms are shown and colored by – log_10_ p-value. **e** ssGSEA scores (y-axis) for selected Reactome pathways across the range of ACTA2 log_2_ intensity (x-axis). Lines are smoothed (Savitzky-Golay, 100 neighbors, 2^nd^ order smoothing) and colored by their Spearman correlation with “Smooth muscle contraction”.

Given the central role of ACTA2 in the phenotypic axis, we examined correlating proteins (Figure 2c). ACTA2 was positively correlated with canonical contractile markers (CNN1 ρ = 0.97 and MYH11 ρ = 0.97), confirming the coherence of the contractile axis. Notably, PDLIM7 (ρ = 0.94) and SORBS1 (ρ = 0.93), involved in adhesion and actin remodeling, ranked among the strongest positive correlators and represent candidates for further functional investigation in the maintenance of the contractile phenotype^22^. Proteins negatively correlated with ACTA2 included established dedifferentiation markers, such as LGALS3 (ρ = –0.61) and ICAM1 (ρ = – 0.62)^23^, alongside less discussed proteins (FERMT3 ρ = –0.86, LSP1 ρ = –0.82, NCEH1 ρ = –0.81 and PGD ρ = –0.84). Over-representation analysis on the top 500 proteins most anti-correlated with ACTA2 revealed enrichment for phagocyte effector machinery, antigen processing and a metabolic shift, recapitulating established effector programs of phenotypic switching^13,14,24^ (Figure 2d).

To confirm that these programs vary continuously with ACTA2 abundance at the individual neighborhood level, we performed single-sample gene set enrichment analysis (ssGSEA) across Reactome pathways (Figure 2e). Smooth muscle contraction was strongly positively correlated with ACTA2 abundance, confirming presence of the contractile phenotype at the single-neighborhood level (Extended Figure 2f). In contrast, lipid metabolism, translation, respiratory electron transport and interferon signaling all increased as ACTA2 decreased. This reinforces the direct relationship between loss of contractile identity and activation of dedifferentiation programs in human atherosclerotic plaques.

### Spatial organization reflects molecular functionalities of VSMCs

We hypothesized that VSMC molecular signatures are shaped by their spatial position within the plaque. To test this systematically, we applied a quantitative spatial scoring system in which each neighborhood was assigned a normalized radial position relative to the luminal center of the plaque, with scores ranging from 0 (center of the lumen) to 1 (plaque periphery)^25^. The score distribution across all VSMC neighborhoods recapitulated the established spatial distribution of VSMCs in plaques: most VSMCs (46%) were localized at the plaque periphery (score 0.8 – 1.0), a smaller population (12%) was adjacent to the lumen (0.0 – 0.2) and the remainder were distributed throughout the mid-plaque region (0.2 – 0.8) (Figure 3a).

**Figure 3:**
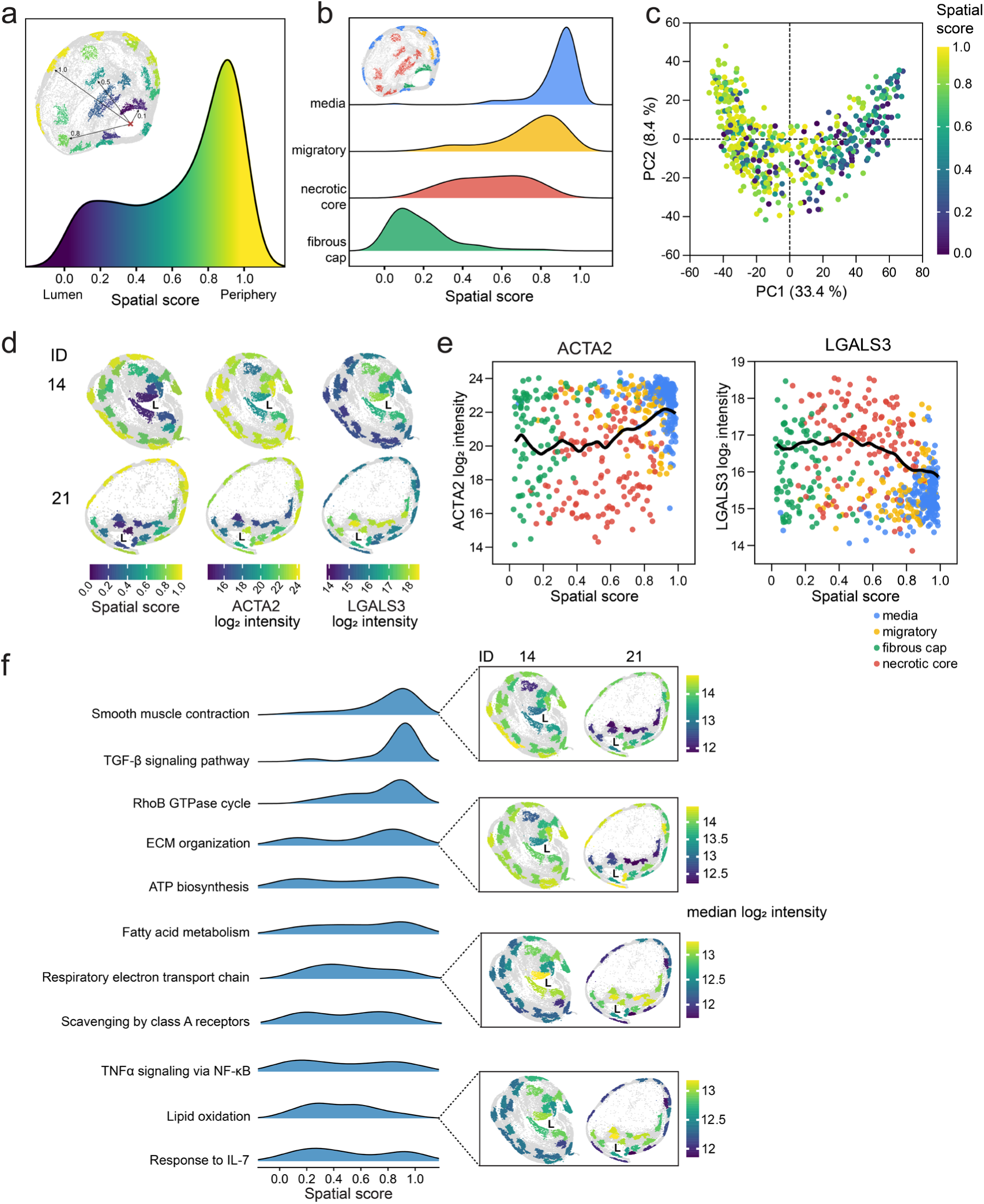
Spatial organization reflects molecular functionalities of VSMCs. **a** Density distribution of spatial scores across all neighborhoods (n = 535). Low scores correspond to neighborhoods near the plaque lumen, high scores to the periphery. **b** Density distributions of spatial scores stratified by histological plaque subregion (media, migratory, fibrous cap, necrotic core). **c** PCA of VSMC neighborhood proteomes (n = 535) colored by spatial score. **d** ACTA2+ shapes colored by spatial score, ACTA2 or LGALS3 log_2_ intensity. Plaques 14 and 21 are shown. L = plaque lumen. **e** ACTA2 and LGALS3 log_2_ intensity versus spatial score. Neighborhoods are colored by histological subregion, black lines indicate LOWESS smoothing. **f** Spatial patterns of selected biological pathways. Left: Density distributions of spatial scores for the 50 neighborhoods with highest median log_2_ intensity for each pathway. Right: ACTA2+ shapes colored by median log_2_ intensity of indicated pathways, plaque 14 and 21 are shown. L = plaque lumen.

To confirm that our scoring system accurately reflects the histopathological architecture of plaques, we mapped spatial scores onto histological plaque subregions. While conventional classification defines three primary subregions: media, necrotic core and fibrous cap^26^, we included an additional ‘migratory’ subregion in our analysis, which we defined as adjacent to the media but oriented toward the plaque interior. Subregions were assigned manually through careful histological examination of IF and HE-stained sections. The resulting distribution matched the expected plaque architecture: media neighborhoods exhibited the highest spatial scores, followed by migratory neighborhoods. Necrotic core neighborhoods had intermediate scores and fibrous cap neighborhoods the lowest scores (Figure 3b). This confirms that our scoring system provides a robust framework for correlating spatial position with molecular signatures.

We then analyzed whether VSMCs display region-specific proteome signatures corresponding to their distinct functional roles. Overlaying the spatial score of each neighborhood onto the PCA plot revealed an enrichment pattern along PC1 that mirrored the ACTA2 gradient (Figure 3c, Spearman ρ = –0.425, p < 0.0001). This shows that spatial context contributes to the proteomic variability and represents a major axis of heterogeneity.

Subsequently, we projected ACTA2 abundance and spatial score onto ACTA2+ shapes. This revealed concordant distribution patterns across most plaques: high ACTA2 intensities co-localized with high spatial scores toward the plaque periphery, while both decreased toward the lumen (Figure 3d, Extended Figure 3a). Across all neighborhoods, ACTA2 intensity correlated modestly but significantly with the spatial score (Pearson r = 0.346, p < 0.0001). LOWESS data smoothing revealed a multimodal pattern in which ACTA2-high neighborhoods clustered toward the periphery (Figure 3e). The established dedifferentiation marker LGALS3^23^, which we had shown to be anti-correlated with ACTA2 (Spearman ρ = –0.61), displayed the opposite spatial pattern: low expression at the periphery and higher expression toward the lumen (Pearson r = –0.317, p < 0.0001).

Finally, we examined how molecular programs enriched in VSMC neighborhoods relate to their spatial organization. We calculated pathway enrichment scores for each neighborhood and analyzed their distribution across five spatial bins (Extended Figures 4a-b). Our analysis revealed distinct spatial patterns for different functional modules. SMC contraction was strongly enriched toward the plaque periphery, ECM organization was enriched both at the plaque periphery and at the plaque lumen, while the lumen and mid-plaque regions were associated with metabolic reprogramming (respiratory electron transport chain, lipid oxidation) (Figure 3f).

These spatial-molecular relationships are consistent with the pathophysiological progression of VSMC phenotypic switching. Contractile VSMCs in the media undergo progressive dedifferentiation as they migrate into the plaque, transitioning from a quiescent, contractile state to metabolically active phenotypes with additional effector functions^5^. Our spatial approach resolves phenotypic switching as a graded, spatially organized continuum of functional states in advanced human atherosclerosis and carotid artery disease.

### Proteomic atlas of primary VSMC phenotype models

To enable functional annotation of the heterogeneous VSMC proteomes observed in plaques, we generated proteomic profiles of key VSMC phenotypes. We exposed patient-derived primary carotid artery SMCs to five stimuli known to induce distinct phenotypes: TGF-β^27^, PDGF-BB^28^, osteogenic stimulants^29^, IL-1β^30^ and free cholesterol^31,32^. For each phenotype, we obtained cellular proteome and secretome by analyzing samples on the Orbitrap Astral mass spectrometer (Figure 4a, Methods).

**Figure 4:**
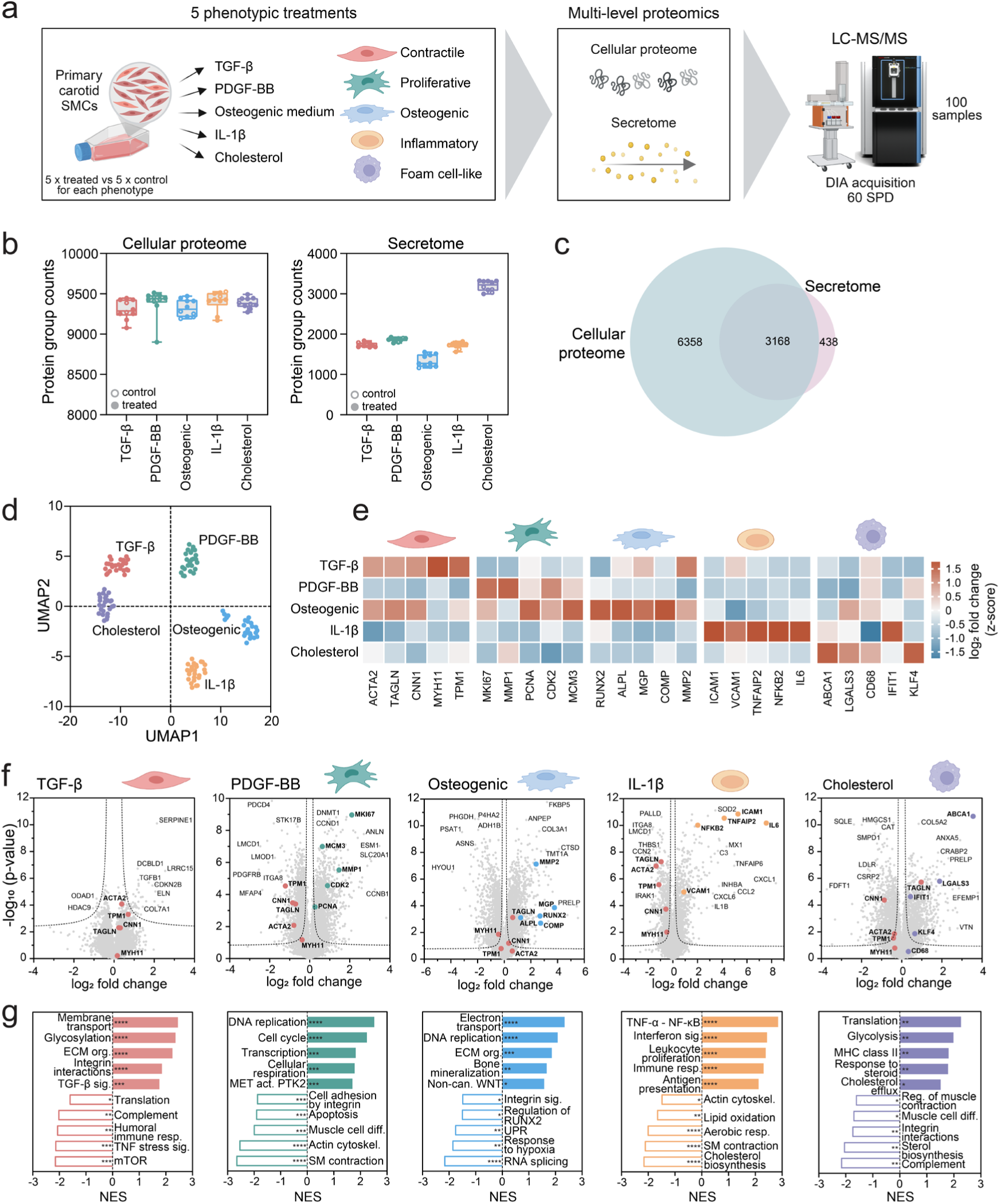
Proteomic atlas of primary VSMC phenotype models. **a** Experimental design of phenotypic treatments. Primary human carotid SMCs were treated with TGF-β, PDGF-BB, osteogenic medium, IL-1β or cholesterol. Cellular proteome and secretome were acquired by LC-MS/MS. **b** Protein group counts per condition for cellular proteome (left) and secretome (right) (n = 5 per condition). **c** Overlap of proteins detected in cellular proteome and secretome at the gene level. **d** UMAP (number of neighbors = 15, number of components = 2, random state = 1, minimum distance = 0.8, metric = Euclidean) of treated-vs-control log_2_ fold changes across all phenotypes at the cellular proteome level. **e** Heatmap of phenotype markers in the cellular proteome. Color indicates z-scored log_2_ fold change versus control. **f** Volcano plots of treated versus control samples for each phenotype (unpaired two-sided Student’s t-test, FDR < 0.05, s_0_ = 0.1). Cellular proteome and secretome data were merged for testing. Selected contractile markers are colored in red, dedifferentiation markers are colored by their respective phenotype. **g** Gene-set enrichment analysis of treated versus control samples for each phenotype using log_2_ fold change as the ranking metric. NES = normalized enrichment score. FDR q-value < 0.001 (****), < 0.05 (***), < 0.1 (**), < 0.25 (*).

Our *in vitro* experiments quantified more than 9,000 proteins per sample in the cellular proteome and up to 3,000 proteins in the secretome (Figure 4b); about 400 proteins were uniquely detected in the secretome (Figure 4c). As expected, the secretome was enriched for secreted and ECM proteins, including metalloproteases and proteins involved in immunity. In contrast, the cellular proteome was enriched for intracellular proteins, including ribosomal proteins, initiation factors and RNA-binding proteins (Extended Figure 5a). This shows that our complementary approach captures the full spectrum of molecular changes, enabling comprehensive characterization of both intracellular signaling mechanisms and extracellular matrix remodeling processes.

To evaluate treatment effects, we calculated log_2_ fold changes between all treatment-control pairs (5 controls, 5 treated) at the cellular proteome level, generating 25 data points per treatment condition. UMAP (Uniform Manifold Approximation and Projection) analysis revealed distinct clustering patterns (Figure 4d), indicating that each treatment induces unique proteome shifts. We then computed Pearson correlation coefficients using log_2_ fold change values across all treatments to identify phenotypic similarities between treatments (Extended Figure 5b). TGF-β and PDGF-BB treatments exhibited the highest correlation (r = 0.37), indicating the greatest phenotypic similarity across all treatment pairs.

To determine whether the observed proteome shifts correlate with expected differences of dedifferentiated phenotypes, we examined a panel of known marker proteins across all treatment conditions (Figure 4e). Contractile markers (e.g. ACTA2, TAGLN, MYH11) showed subtle upregulation in TGF-β treated samples. In contrast, these markers were consistently downregulated in PDGF-BB, IL-1β and cholesterol treated samples, while the osteogenic treatment displayed a divergent pattern. At the same time, markers of dedifferentiated states were selectively upregulated in their respective treatment conditions. Comprehensive analysis of the proteomes across all treatments provided further evidence of unique proteome alterations consistent with predicted phenotypic transitions (Extended Figure 5c).

We merged cellular proteome and secretome datasets to enable joint downstream analysis of intracellular and extracellular protein changes, retaining the entry with the largest absolute log_2_ fold change across treatments for proteins detected in both datasets, allowing us to capture the most pronounced changes in protein abundance across the entire system. Pair-wise comparisons across treatment-control for all phenotypes revealed alterations induced by each treatment (Figure 4f). TGF-β induced the fewest changes, suggesting that untreated cells in culture most closely resemble this phenotype, maintaining baseline contractility. Conversely, PDGF-BB and osteogenic treatments generated the largest number of significantly changing proteins, followed by cholesterol and IL-1β treatment (Extended Figure 5d).

Gene set enrichment analysis (GSEA) revealed pathways associated with phenotypic transitions (Figure 4g). TGF-β treatment induced subtle changes, pushing cells toward a modest contractile-matrix-producing state, with reduced stress signaling. PDGF-BB induced a proliferative state, characterized by upregulated DNA replication and cell cycle proteins, while simultaneously downregulating smooth muscle contraction. Osteogenic stimuli induced biomineralization and ECM remodeling alongside a strong proliferative signature and upregulated oxidative phosphorylation. IL-1β treatment downregulated smooth muscle contraction and oxidative metabolism, accompanied by activation of inflammatory signaling pathways. Cholesterol reduced muscle-associated programs alongside shutdown of sterol biosynthesis, while activating translation, glycolysis and MHC II– and IFN-associated programs.

Analysis of upstream regulators further validated our treatment results, identifying TGF-β, the PDGF complex, all three osteogenic stimulants (ascorbic acid shown as example), IL-1β and cholesterol as significantly activated upstream regulators for their respective treatments (Extended Figure 6a-e). For clarity, from here on, we refer to each phenotype by its dominant functional identity: contractile (TGF-β), proliferative (PDGF-BB), inflammatory (IL-1β), osteogenic, and foam cell-like (cholesterol).

This proteomic atlas of VSMC phenotypes provides a reference for understanding phenotypic switching and a resource for future studies in vascular biology and atherosclerosis.

### Deep learning framework identifies association of phenotypic switching and plaque instability

A central challenge in spatial proteomics is linking *in situ* molecular signatures to defined functional states. To address this, we leveraged our primary VSMC phenotype proteomes as a functional reference for the DVP-derived spatial neighborhood data. However, integration of tissue and cell-culture proteomes is challenging due to systematic differences in proteome depth, dynamic range, and technical batch effects. To address this, we implemented a semi-supervised conditional variational autoencoder (CVAE) deep learning model (Figure 5a). CVAE consists of three components (encoder, decoder, phenotype discriminator) and we optimized it with a composite loss that enforces accurate reconstruction, latent-space regularization, phenotype discriminability and sparsity. By conditioning the latent embedding on well-characterized *in vitro* phenotypes, CVAE projects spatial proteomes onto a continuous phenotype manifold, thereby enabling functional annotation of spatially defined molecular patterns.

**Figure 5:**
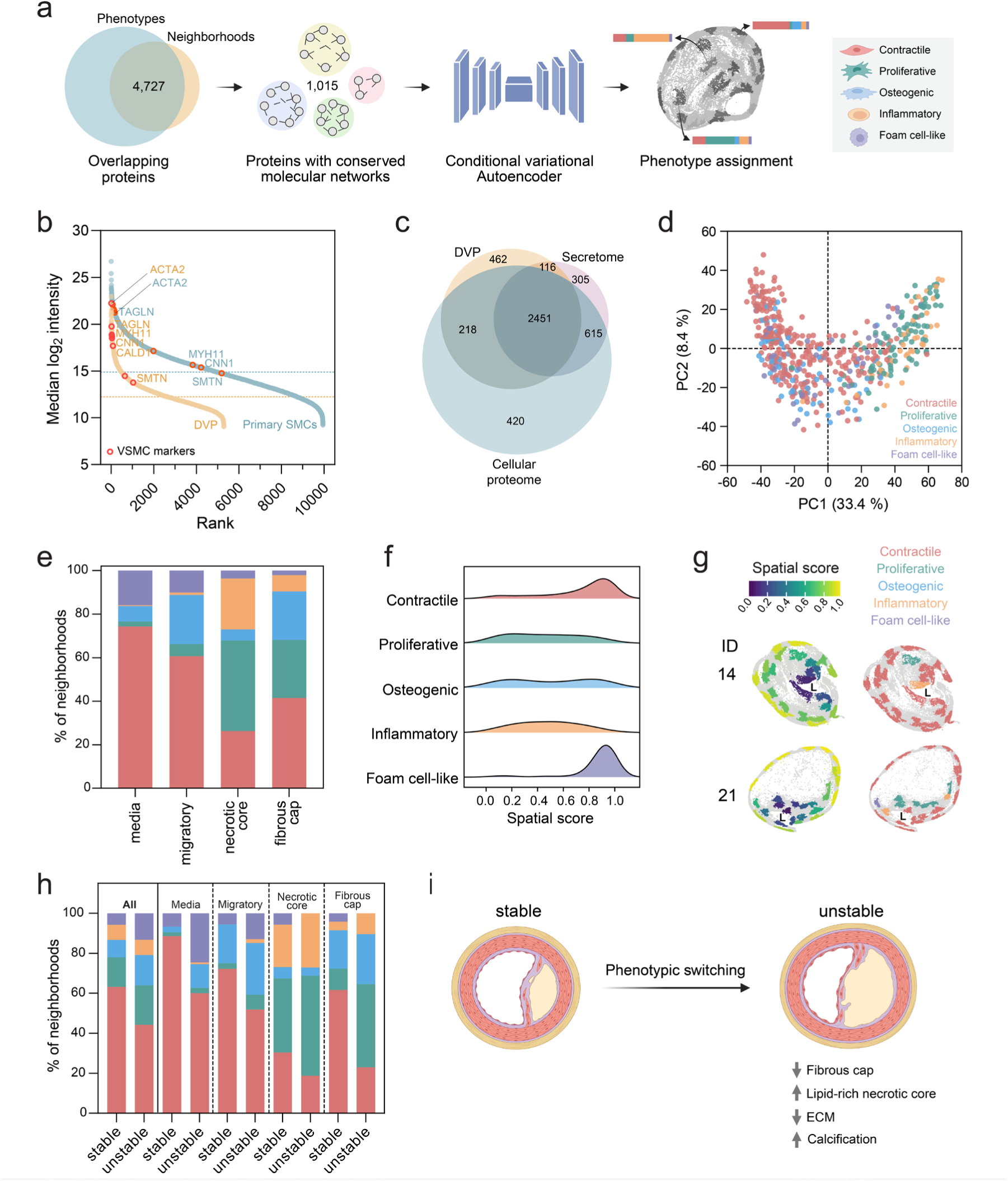
Deep learning framework identifies association of phenotypic switching and plaque instability. Each neighborhood is represented by the phenotype with the highest probability. **a** Computational framework integrating spatial neighborhood proteomes with primary VSMC phenotype references. Proteins detected in both datasets (n = 4,727) were selected and proteins with conserved molecular networks (n = 1,015) were used as input for the conditional variational autoencoder. A random forest classifier was used to assign phenotype probabilities to each neighborhood. **b** Rank plots of median log_2_ intensity for the DVP proteome (orange) and the primary VSMC cellular proteome (blue). Hallmark VSMC markers are highlighted in red. **c** Overlap of proteins detected in the DVP proteome, primary VSMC cellular proteome and secretome at the gene level. **d** PCA of VSMC neighborhood proteomes (n = 535), colored by assigned phenotype. **e** Phenotype proportions across histological plaque subregions. **f** Density distribution of spatial scores for each phenotype. **g** ACTA2+ shapes colored by spatial score (left) and phenotype identity (right), shown for plaques 14 and 21. L = plaque lumen. **h** Phenotype proportions within histological plaque subregions reflecting neighborhood-level counts, stratified by plaque stability. **i** Proposed link between phenotypic switching and plaque instability.

Cell culture-derived reference proteomes comprised 9,948 protein groups, whereas tissue samples yielded 5,287 protein groups (Figure 5b), reflecting the complexity of FFPE-based workflows relative to cell culture conditions. Importantly, 4,727 protein groups were shared between both datasets (89% overlap of the tissue proteome) (Figure 5c), providing a robust core for phenotype inference. To focus on functionally relevant proteins and minimize dataset-specific noise, we constructed molecular networks and retained only 1,015 protein groups whose network topology was conserved across both datasets (Spearman ρ > 0.3). This filtered protein set served as the input for CVAE, which embedded each neighborhood proteome into a phenotype-conditioned latent space, followed by a random forest classifier that calculated phenotype probabilities for each neighborhood.

UMAP visualization of the resulting latent embedding of the VSMC neighborhoods confirmed that CVAE successfully separated the five reference phenotypes (Extended Figure 7a), indicating that the model learned phenotype-discriminative features from the input proteomes. Across all 535 neighborhoods, CVAE returned probability scores distributed across all five reference phenotypes (Extended Figure 7b), reflecting the multi-cellular composition of each neighborhood, where collected cells may simultaneously occupy different phenotypic states. For all subsequent analyses we used the highest-probability phenotype as the per-neighborhood assignment. We confirmed that contractile (TGF-β) probability tracked ACTA2 intensity (Extended Figure 7c) and that assignment matched phenotype-specific marker proteins, showing that contractile proteins were reduced in dedifferentiated states, while dedifferentiation markers were upregulated (Extended Figure 7d).

To verify that CVAE preserves the dominant biological structures of the DVP data, we projected the phenotypes with the highest probability for each neighborhood onto the PCA plot of the DVP data (Figure 5d). Contractile phenotypes localized predominantly at low PC1 values whereas dedifferentiated phenotypes mapped to higher PC1 values. Hence, our CVAE-based integrative analysis recapitulated the continuum defined by ACTA2 (see Figure 2a).

Spatial mapping of neighborhood phenotypes across plaque subregions revealed patterns that align with known plaque biology (Figure 5e). The media was dominated by contractile VSMCs, consistent with their role in maintaining vessel wall integrity. Dedifferentiated states (proliferative, osteogenic, inflammatory) concentrated within necrotic cores and fibrous caps. Interestingly, foam cell-like neighborhoods localized preferentially toward the media and migratory zone. This unexpected distribution may reflect early exposure to accumulating lipids, analogous to the cholesterol loading conditions in our *in vitro* atlas (Figure 4e-g). Spatial score distributions for each phenotype confirmed the structured organization of VSMC phenotypes in human plaques (Figure 5f), which was reproducible at the level of individual plaques (Figure 5g, Extended Figure 7e).

Having established a robust framework for phenotype mapping, we next examined how VSMC compositions differed between histologically defined stable and unstable plaques. Unstable plaques showed a reduction in the proportion of neighborhoods classified as contractile (–20.7%, p-value = 0.037), accompanied by a corresponding increase of dedifferentiated states (Figure 5h). This shift was consistent across patients and was not driven by individual outliers (Extended Figure 7f). Analysis across histological subregions revealed that loss of contractile cells was most pronounced in the media (–25.5%, p-value = 0.044) and fibrous cap (–33.9%, p-value = 0.024), accompanied by an expansion of proliferative cells in the fibrous cap (+27.5%, p-value = 0.036) (Figure 5h). To exclude potential sampling bias, we confirmed that subregion proportions were comparable between both stability groups (Extended Figures 7g-h). Overall, these shifts link VSMC phenotypic switching to plaque instability (Figure 5i). Given that dedifferentiated states were enriched in unstable plaques, we next sought to identify and validate candidate proteins associated with these phenotypic transitions.

### Perturbation experiments in primary VSMCs confirm functional associations

To nominate candidate proteins for functional validation, we leveraged both our DVP and cell culture dataset. We focused on proteins upregulated in dedifferentiated states in the primary VSMC atlas (cellular proteome) that showed concordant enrichment in the corresponding CVAE-assigned plaque neighborhoods, relative to contractile-assigned neighborhoods. From this set, we selected candidates with plausible roles in VSMC phenotypic switching that span the architecture of the atlas: one pan-phenotypic protein (TNC), upregulated across all dedifferentiated phenotypes, and three phenotype-specific proteins (PLK1: proliferative, AEBP1: osteogenic, TNFAIP2: inflammatory), each predominantly upregulated in a single phenotype (Figure 6a, Extended Figure 8a). Additional candidate proteins not pursued here are listed in Extended Data Table 2.

**Figure 6:**
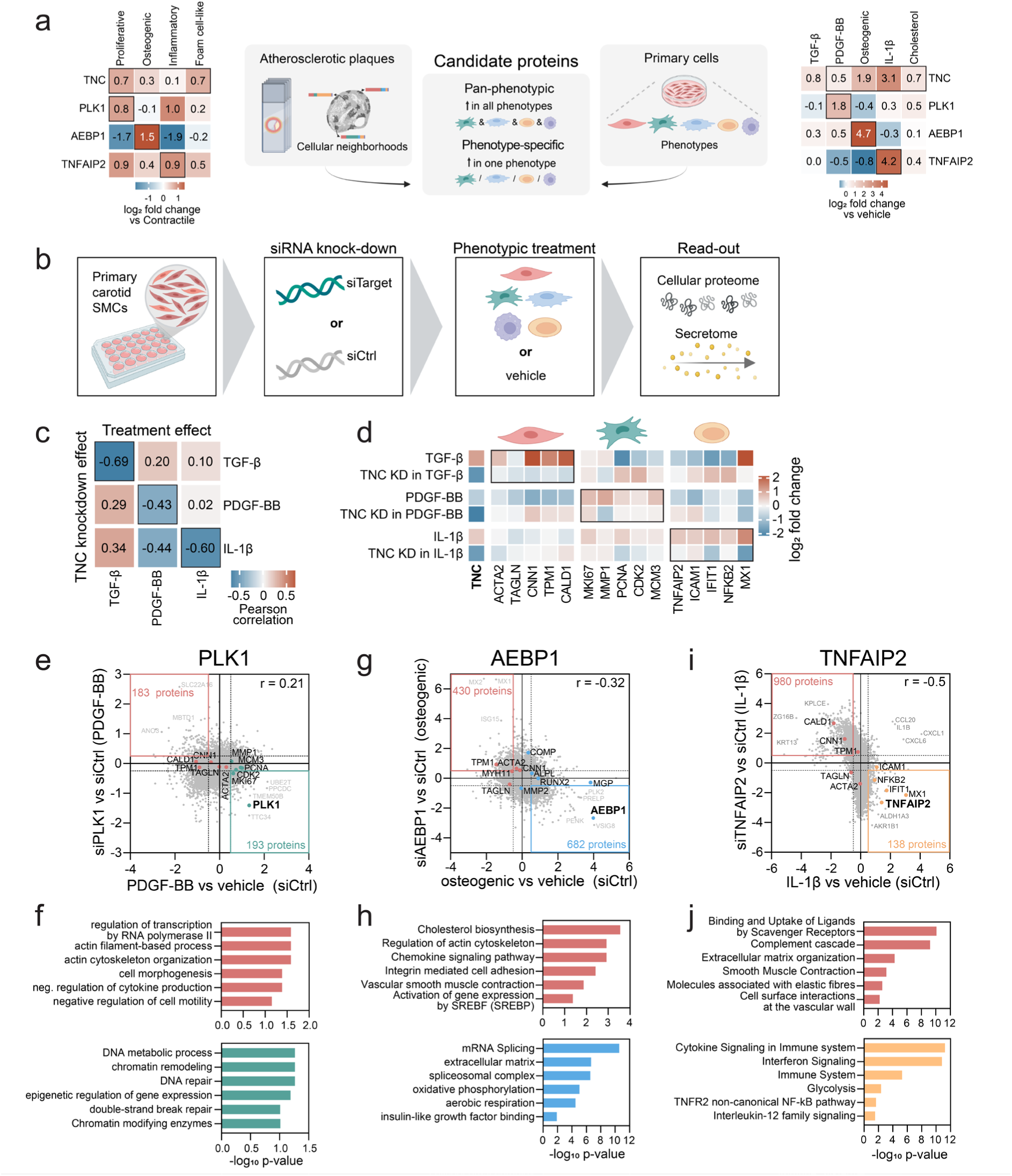
Perturbation experiments in primary VSMCs confirm functional associations. **a** Selection of candidate proteins. Expression patterns of TNC, PLK1, AEBP1 and TNFAIP2 in primary VSMC reference data (right, treated vs vehicle) and DVP cohort (each dedifferentiated vs contractile-assigned neighborhoods). **b** Design of perturbation experiments. Primary human carotid SMCs were transfected with siRNA against the candidate target (or scrambled siCtrl), exposed to phenotypic treatments and profiled by LC-MS/MS. **c** Concordance between treatment and TNC-knockdown effects. Pearson correlation of log_2_ fold changes between treatment effect (treatment vs. vehicle under siCtrl) and TNC knockdown effect (siTNC vs. siCtrl under treatment). **d** Heatmap of phenotype markers in the cellular proteome. Color indicates log_2_ fold change versus the corresponding control (vehicle for treatments and siCtrl for KD). **e** Scatter plot of PLK1 knockdown effect versus PDGF-BB treatment effect. **f** Over-representation analysis of proteins (red box) down-regulated by PDGF-BB (log_2_ fold change < –0.5) and rescued by PLK1 knockdown (log_2_ fold change > 0.25); and proteins (green box) up-regulated by PDGF-BB (log_2_ fold change > 0.5) and attenuated by PLK1 knockdown (log_2_ fold change < –0.25). The background for ORA was restricted to PDGF-BB responsive proteins (log_2_ fold change < –0.5 or > 0.5), and an FDR of 0.1 was applied. **g** Scatter plot of AEBP1 knockdown effect versus osteogenic treatment effect. **h** Over-representation analysis of proteins (red box) down-regulated by osteogenic treatment (log_2_ fold change < –0.5) and rescued by AEBP1 knockdown (log_2_ fold change > 0.5), and proteins (blue box) up-regulated by osteogenic treatment (log_2_ fold change > 0.5) and attenuated by AEBP1 knockdown (log_2_ fold change < –0.5). **i** Scatter plot displaying TNFAIP2 knockdown effect versus IL-1β treatment effect. **j** Over-representation analysis of proteins (red box) down-regulated by IL-1β (log_2_ fold change < –0.5) and rescued by TNFAIP2 knockdown (log_2_ fold change > 0. 5), and proteins (orange box) up-regulated by IL-1β (log_2_ fold change > 0.5) and attenuated by TNFAIP2 knockdown (log_2_ fold change < –0. 5).

We performed siRNA-mediated knockdown (KD) of each candidate protein in primary carotid artery VSMCs, followed by phenotype-inducing treatment and measured changes in the cellular proteome and secretome (Figure 6b). We quantified between 8,000 and 10,000 protein groups in the cellular proteome (Extended Figure 8b), with separation of conditions on the PCA level (Extended Figure 8c). Knockdown of each candidate was confirmed at the protein level for all four proteins in cellular proteome (Extended Figure 8d) and for TNC and AEBP1 in the secretome (Extended Figure 8e). Reductions were most pronounced in treatment-induced conditions, consistent with the larger dynamic range available when target expression is elevated by phenotypic stimulation.

Tenascin-C (TNC) was consistently upregulated across all dedifferentiated phenotypes in our phenotype atlas (Figure 6a), making it a prime pan-phenotypic candidate. It is a large matricellular glycoprotein that is upregulated in atherosclerotic plaques and acts as a context-dependent regulator of proliferation, migration, differentiation, and remodeling^33^. To assess whether TNC is necessary for induction of phenotypic programs, we performed siRNA-mediated TNC knockdown prior to TGF-β, PDGF-BB or IL-1β treatment. We compared the proteome-wide effect of TNC knockdown within each treatment context (siTNC vs siCtrl, in each treatment) to the corresponding treatment effect in control cells (treatment vs vehicle in siCtrl cells). We observed strong negative correlations for all three treatments (Figure 6c), indicating that TNC reduction prevents treatment-induced changes. Consistent with this, phenotype-specific marker proteins were induced by each treatment but consistently dampened upon TNC reduction (Figure 6d). For PDGF-BB and IL-1β, downregulation of contractile proteins was likewise attenuated. These results establish TNC as a pan-phenotypic protein necessary for VSMC phenotypic switching in response to TGF-β, PDGF-BB and IL-1β.

Polo-like kinase 1 (PLK1) was selectively upregulated by PDGF-BB in our phenotype atlas (Figure 6a). PLK1 is a serine/threonine kinase required for G2/M progression and is induced in proliferating VSMCs at sites of vascular injury^34^. In contrast to the strong anti-correlation observed for TNC, the proteome-wide effect of PLK1 knockdown in PDGF-BB-treated cells showed weak positive correlation with the PDGF-BB treatment effect (Pearson r = 0.21; Figure 6e). This indicates that PLK1 depletion does not oppose the proliferative switch but instead acts on a subset of the PDGF-BB response. Consistent with a focused mitotic role, proliferation markers (MKI67, CDK2, PCNA, MCM3) were dampened upon PLK1 knockdown. Over-representation analysis (Reactome, GO) revealed that PLK1 depletion attenuated DNA metabolic process, chromatin remodeling and DNA repair programs, while concurrently preserving actin cytoskeleton organization and cell morphogenesis (Figure 6f). These results position PLK1 as a focused effector of the mitotic execution arm; targeting PLK1 could therefore restrain proliferative VSMC expansion in plaques.

Adipocyte enhancer-binding protein 1 (AEBP1) was strongly upregulated by osteogenic treatment in primary VSMCs and in osteogenic-classified VSMC neighborhoods (Figure 6a). AEBP1 is a multifunctional collagen-binding ECM protein and transcriptional repressor with established roles in adipogenesis, ECM organization and VSMC differentiation^35^. The effect of AEBP1 knockdown in osteogenic-treated cells anti-correlated with the osteogenic treatment effect (Pearson r = –0.32) (Figure 6g). Over-representation analysis in Reactome, KEGG and GO confirmed that AEBP1 depletion attenuated osteogenic-induced programs, including ECM organization and oxidative phosphorylation, while preserving contractile and cytoskeletal programs (Figure 6h). This dual signature suggests that AEBP1 contributes to the execution of the osteogenic switching program in VSMCs. In plaques, where osteogenic VSMCs contribute to calcification, AEBP1 inhibition could dampen pathological matrix and calcification programs and favor retention of contractile features.

Tumor necrosis factor-α-induced protein-2 (TNFAIP2) has been linked to atherogenesis through its enhancement of oxidative-stress-driven inflammation^36^, but its role in VSMC-driven processes in atherosclerosis remains largely uncharacterized. In our reference atlas, TNFAIP2 was highly upregulated by IL-1β in primary VSMCs and showed corresponding enrichment in inflammatory-classified VSMC neighborhoods (Figure 6a), suggesting an unappreciated function in inflammatory VSMCs. The proteome-wide effect of TNFAIP2 knockdown in IL-1β-treated cells anti-correlated with the IL-1β-treatment effect (Pearson r = –0.5) (Figure 6i). Over-representation analysis (Reactome) showed that TNFAIP2 depletion attenuated IL-1β-induced inflammatory programs, including cytokine and interferon signaling, while concurrently restoring a subset of contractile and ECM-organization programs (Figure 6j). Given that VSMC-derived inflammatory signaling is a driver of plaque progression and instability^5^, TNFAIP2 inhibition could blunt VSMC participation in the local inflammatory milieu.

Collectively, our phenotype-resolved knockdown experiments confirm TNC as a pan-phenotypic protein and TNFAIP2, AEBP1, and PLK1 as proteins involved in phenotype-specific programs. These molecular switches represent candidate nodes through which normal contractile VSMCs are transformed into pathological phenotypes, pointing to phenotype-specific mechanisms of VSMC dysfunction associated with plaque destabilization and offering potential therapeutic entry points for intervention in human atherosclerosis.

## Discussion

In this study, we present a cell type– and phenotype-resolved spatial proteomic atlas of human atherosclerotic plaques and introduce a computational framework that bridges heterogeneous tissue proteomes and defined functional states. Using DVP, we mapped over 500 VSMC neighborhoods from 24 human carotid plaques and resolved VSMC phenotypic switching on the spatial proteome level. A key methodological contribution is the integration of spatial plaque proteomes with reference profiles from primary VSMC phenotype models through a conditional variational autoencoder. This enabled functional annotation of *in situ* molecular signatures that would otherwise remain uninterpretable. Phenotype-resolved siRNA validation of four targets closes the loop from spatial observation to molecular validation, showing that our framework not only describes VSMC heterogeneity, but can nominate candidate drivers of phenotypic switching and validate them at the proteome level in a phenotype-resolved manner. This turns spatial proteomic observations into molecularly validated and translationally relevant targets in atherosclerosis and vascular disease.

Our findings support a view of VSMCs as active, spatially organized regulators of atherosclerotic plaques. The proteome heterogeneity we observe complements single-cell and spatial transcriptomic studies^13,14,17^ by resolving spatial organization within intact plaque architecture and providing protein-level evidence of functional states. Our data shows that protein-level programs are spatially organized: contractile VSMCs concentrate in the media, while dedifferentiated states populate the necrotic core and fibrous cap. Interestingly, unstable plaques showed a decrease in contractile VSMCs with corresponding expansion of dedifferentiated states, most strikingly in the fibrous cap. This reinforces that VSMC phenotypic switching contributes to plaque destabilization rather than acting as a bystander process, and complements the recent subregion-resolved proteomic signatures of carotid plaque instability^17^ by adding cell type and phenotype resolution.

Functional perturbation of four candidate proteins establishes that the framework moves from observation to application. TNC emerged as a pan-phenotypic protein necessary for the full TGF-β, PDGF-BB and IL-1β responses; TNFAIP2 as a major contributor to the IL-1β inflammatory program; AEBP1 as a contributor to osteogenic remodeling, and PLK1 as a focal effector of the mitotic arm of the proliferative response. Each defines a distinct therapeutic entry point and supports a growing view of VSMC phenotypic switching as a tractable therapeutic target in human atherosclerosis^37^. The four validated candidates could be further tested in human plaque *ex vivo* culture models^38,39^ and advanced atherosclerosis-relevant *in vivo* systems^40^ to assess whether their modulation can stabilize vulnerable lesions. While the concordance between *in vitro* phenotypes and *in situ* spatial proteomic signatures provides initial evidence, these models will be essential to establish whether identified functions translate into complex and multicellular environments.

Several limitations should be considered. Most fundamentally, the cohort size of 24 plaques limits the generalizability of clinical observations, and external validation in larger cohorts will be important to consolidate our results, especially the exploratory phenotype differences between stable and unstable plaques. Another consideration concerns the inherent complexity of tissue-derived proteomes, which are susceptible to signals from surrounding material. Our DVP approach directly addresses this through targeted laser microdissection of immunofluorescence defined cell populations, enriching for cell type-specific proteomes and reducing contributions from surrounding material. Residual background signals are further mitigated by our CVAE, which anchors tissue signatures against reference proteomes derived from pure primary VSMC populations, thereby isolating cell type-specific signals from the complex plaque tissue milieu.

A third limitation concerns our reliance on ACTA2 as the primary marker for VSMC identification. While ACTA2 is widely used to identify VSMCs in vascular tissue, it is not strictly VSMC-exclusive and can be expressed in other stromal cell types, most notably myofibroblasts and pericytes^7^. Definitive attribution of ACTA2+ cells to the VSMC lineage would ultimately require lineage-tracing approaches^41^, which are not feasible in human tissue. However, in our dataset, ACTA2 was detected in all 535 profiled neighborhoods, ranking in the top 10% of protein intensities across the dataset. In addition, other canonical VSMC markers, including MYH11, CNN1 and TAGLN were equally ubiquitous, confirming that the dissected material is enriched for VSMC proteomes. Finally, the five *in vitro* phenotype stimulation models might not span the full range of VSMC states observed in human plaques but were chosen to capture the dominant axes of switching reported in the literature^5,14^. CVAE itself is framed as an interpretable integration tool rather than a phenotype-assignment ground truth. Its output probabilities reflect the relative similarity of each neighborhood to each reference state rather than discrete phenotype calls.

Our work links spatially resolved proteomes to defined functional cell states, closing a gap that has limited the interpretation of spatial tissue proteomics. Its application to atherosclerotic plaques mapped VSMC plasticity at the protein level, established phenotypic switching as a key feature of plaque instability, and nominated candidate proteins as potential therapeutic targets. This framework, combining cell type-resolved spatial proteomics with a controlled reference atlas and integrative deep learning, is directly applicable to any tissue context in which heterogeneous cellular states must be assigned functional meaning, including cancer microenvironments, fibrotic tissues, and developmental contexts in which cell plasticity shapes outcome.

## Methods

### Human carotid plaque collection

Carotid plaques were collected from patients undergoing carotid endarterectomy, sourced from the Munich Vascular Biobank. Informed written consent for biobank sample contributions was obtained from all participants in accordance with the principles of the Declaration of Helsinki. The study received approval from the local ethics committee of the Technical University of Munich (approval numbers: 2799/10 and 2023-297) and clinical data was retrieved from anonymized electronic patient records.

After surgical retrieval, samples were transferred into RNAlater to preserve RNA during transport and subsequent processing. Plaques were fixed with 4% paraformaldehyde (PFA) for 24 hours and decalcified (Entkalker soft SOLVAGREEN) for five days. Following decalcification, plaques were embedded in paraffin and stored at room temperature as formalin-fixed paraffin-embedded (FFPE) blocks.

### Cohort design

24 plaques were selected based on the histological availability of the media layer and abundant cellular content, especially VSMC presence, based on HE– and EvG stained tissue sections, along with availability of comprehensive clinical data. In addition, cases were selected to maintain a covariate balance between stable and unstable patient subgroups (Extended Data Table 1).

### DVP workflow

*Immunofluorescence Staining –* Polyethylene naphthalate (PEN) membrane slides (2 µm, MicroDissect GmbH MDG3P40AK) were coated with 25 µl Chrome Alum-Gelatin Adhesive (Newcomer Supply 1033A) and dried at room temperature overnight. 3 µm thick sections of each FFPE plaque sample were mounted on membrane slides. Prior to immunofluorescence staining, slides were heated at 55 °C for 30 min. Then, slides were deparaffinized and rehydrated (2 x 2 min Xylol, 2 x 1 min 100% ethanol, 2 x 1 min 90% ethanol, 2 x 1 min 75% ethanol, 2 x 1 min 30% ethanol, 2 x 1 min ddH_2_O). Antigen retrieval was performed using glycerol-supplemented Antigen Retrieval buffer (DAKO pH9 S2367 + 10% Glycerol) in a water bath at 88 °C for 20 min, followed by a cooldown at room temperature for 20 min. Slides were washed 2 x 3 min in water and blocked with 5% BSA in PBS + 0.5% Triton X-100 for 1 h. The primary antibody (ACTA2 ab7817 1:200, CD68 ab213363 1:100, CD3 DAKO A0452 1:100) was incubated in Renoir Red Diluent (Biocare Medical, PD904M) overnight at 4 °C in a humid staining chamber. After washing 3 x 3 min in PBS, secondary antibodies (Alexa anti-mouse IgG 750 Cy7 A21039 1:400, Alexa anti-rabbit 647 Cy5 A21242 1:400) were incubated in 1% BSA in PBS for 1 h at room temperature. Slides were washed 3 x 3 min in PBS and counterstained with Sytox Green (Invitrogen 57020 1:40,000 in water) for 10 min. Slides were washed again 3 x 3 min in water. After membrane puncture, sealing and edging of calibration crosses, tissue sections were coverslipped using Slowfade Diamond Antifade Mountant (Invitrogen S36963). Imaging was performed using a Zeiss Axioscan system at 20 x magnification (Plan-Apochromat 20x/0.8 M27 objective). Illumination time was adapted for each channel. Stitching was performed offline using the Zeiss Zen Imaging software.

*Segmentation & Clustering* – Images were imported as.czi files into BIAS (Biological Image Analysis Software) and re-tiled using 2048 x 2048 pixels. Resulting image tiles were used for threshold-based segmentation in QuPath^42^ using the positive cell detection feature with the following settings: requested pixel size: 0.173 µm, background radius: 15 µm, median filter radius: 0 µm, sigma: 1.8 µm, area range: 30-1000 µm^2^, split by shape, cell expansion: 1, smooth boundaries. The intensity threshold was adapted for all 24 plaques individually, using the respective negative stained control as reference. Resulting contours were re-imported into BIAS and exported together with three calibration crosses. Contour outlines were simplified by removing 90% of data points. To prevent membrane instability while cutting multiple adjacent shapes, every 4th shape was randomly deleted.

Clustering of ACTA2+ shapes into neighborhoods was performed using a custom-made clustering algorithm. First, all shapes were converted to polygons. Then, user-defined centroids were used to assign cluster identifiers to shapes using a custom implementation of the k-means algorithm. In a second clustering step, HDBSCAN (hierarchical density-based spatial clustering of applications with noise) was used to find clusters of shapes inside each hard cluster from primary clustering. This ensures that only the most densely associated shapes are selected to represent the cluster. The number of clusters (= neighborhoods) was adapted to plaque size and ACTA2+ area for each plaque (Extended Figure 1e). ACTA2+ area was determined using a pixel classifier in QuPath. 400 shapes per neighborhood were chosen, resulting in a median area of 28,500 µm^2^ per neighborhood. Contours were exported as.xml files.

*Laser microdissection* – Resulting contour outlines for each neighborhood were laser microdissected using an LMD7 (Leica) with a 63x objective. First, calibration crosses were aligned. Then, all 400 shapes of each neighborhood were dissected into 384-well plates using the following middle pulse settings: power: 55, aperture: 1, speed: 20, middle pulse count: 1, final pulse: 1, head current: 60%, pulse frequency: 3009, offset: 190. 384-well plates were sealed, centrifuged at 1,000 g for 5 min and frozen at –20 °C until sample preparation.

*Sample preparation* – Sample preparation was performed using a Bravo pipetting robot (Agilent). First, wells were washed with 4 x 7 µl 100% acetonitrile from each side and dried in a SpeedVac at 45 °C for 20 min. Then, 6 µl of 60 mM TEAB (Sigma-Aldrich T7408) with 0.013% DDM (Sigma-Aldrich D5172) in ddH_2_O was added to each well. The plate was sealed with two adhesive foils and heated for 60 min at 95 °C in a 384-well thermal cycler (Eppendorf). 1 µl of 80% acetonitrile in 60 mM TEAB was added and heated again for 60 min at 75 °C. For digestion, 1 µl with 4 ng Trypsin (Sigma-Aldrich) and 6 ng LysC (Wako) were added and incubated overnight at 37 °C. After 16 h, the reaction was quenched by adding TFA to a final concentration of 1%. Samples were stored at –20 °C until acquisition.

*Peptide loading* – Prior to LC-MS/MS acquisition, samples were loaded onto C18 Evotips (Evotip Pure, EvoSep). Evotips were activated in 1-propanol, washed with 70 µl buffer B (0.1% formic acid in acetonitrile) and centrifuged at 700 g for 1 minute. After re-activation in 1-propanol, tips were washed with 70 µl buffer A (0.1% formic acid). Prior to sample transfer, 100 µl buffer A were added followed by a quick spin-down. After initial sample transfer, each well was rinsed with 10 µl buffer A and also loaded. Sample binding was performed by centrifugation at 500 g for 3 minutes. Tips were washed with 70 µl buffer A and overlaid with 120 µl buffer A. Loaded tips were stored at 4 °C prior to LC-MS/MS acquisition.

*Mass spectrometry acquisition* – Peptide samples were separated by the Evosep One LC system using the whisper zoom 80 SPD gradient connected to an Orbitrap Astral (Thermo). An Aurora Rapid column of 5 cm length, 75 µm internal diameter, packed with 1.7 µm C18 beads (IonOpticks AUR4-5075C18) was used (50 °C). Electrospray voltage was set to 1,900 V and a FAIMS Pro interface with compensation voltage of –40 V was used. Data-independent acquisition was performed over a scan range of 380-980 m/z with 100 variable-width windows and 11 ms maximum injection time.

### Primary VSMCs workflow

*Treatments* – Primary cell cultures of human carotid artery smooth muscle cells (PB-3514-05a-PeloBiotech) at passage 6 were used for all experiments. Cells were grown to 70% confluency in complete human smooth muscle cell growth medium (PeloBiotech). Prior to treatment, cells were starved. All treatments were performed in replicates of five with their respective vehicle controls in T-75 flasks. The table below displays all stimuli and respective conditions used:

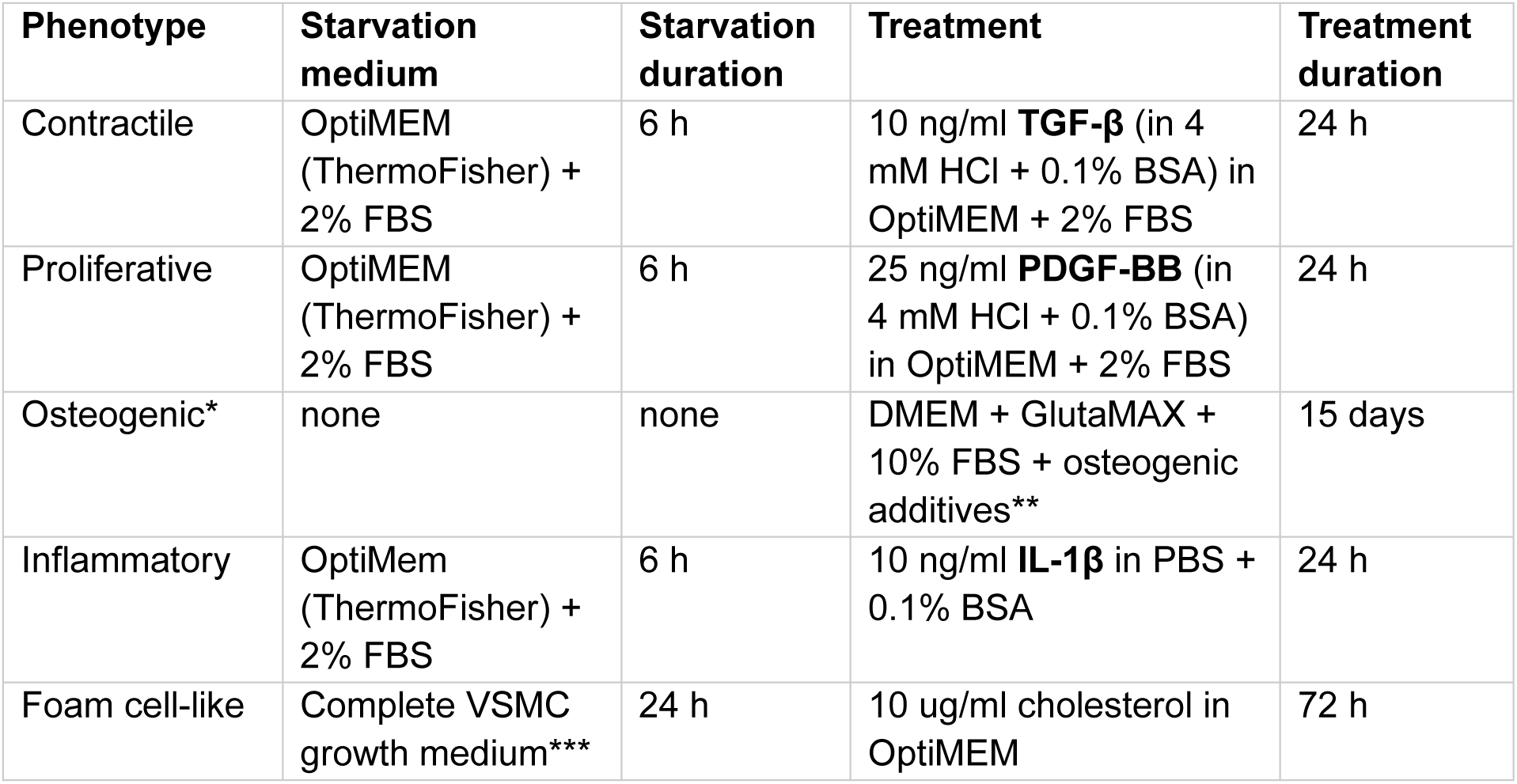
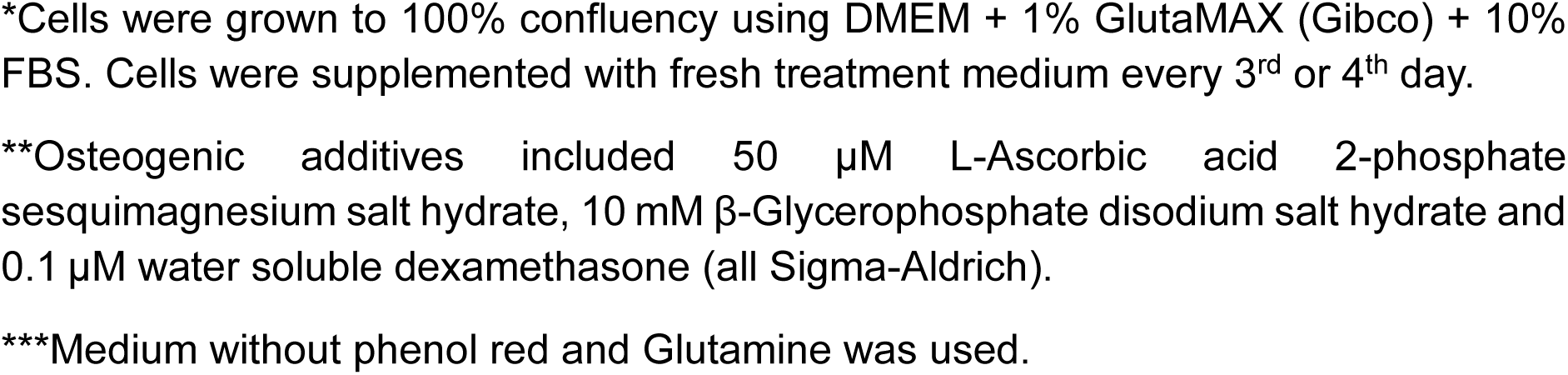

| Phenotype | Starvation medium | Starvation duration | Treatment | Treatment duration |
| --- | --- | --- | --- | --- |
| Contractile | OptiMEM (ThermoFisher) + 2% FBS | 6 h | 10 ng/ml **TGF-β** (in 4 mM HCl + 0.1% BSA) in OptiMEM + 2% FBS | 24 h |
| Proliferative | OptiMEM (ThermoFisher) + 2% FBS | 6 h | 25 ng/ml **PDGF-BB** (in 4 mM HCl + 0.1% BSA) in OptiMEM + 2% FBS | 24 h |
| Osteogenic* | none | none | DMEM + GlutaMAX + 10% FBS + osteogenic additives** | 15 days |
| Inflammatory | OptiMem (ThermoFisher) + 2% FBS | 6 h | 10 ng/ml **IL-1β** in PBS + 0.1% BSA | 24 h |
| Foam cell-like | Complete VSMC growth medium*** | 24 h | 10 ug/ml cholesterol in OptiMEM | 72 h |

\*Cells were grown to 100% confluency using DMEM + 1% GlutaMAX (Gibco) + 10% FBS. Cells were supplemented with fresh treatment medium every 3<sup>rd</sup> or 4<sup>th</sup> day.
\*\*Osteogenic additives included 50 $\mu$ M L-Ascorbic acid 2-phosphate sesquimagnesium salt hydrate, 10 mM $\beta$ -Glycerophosphate disodium salt hydrate and 0.1 $\mu$ M water soluble dexamethasone (all Sigma-Aldrich).
\*\*\*Medium without phenol red and Glutamine was used.

*siRNA-mediated knockdowns* – All knockdown experiments were performed with respective vehicle controls in 24-well plates. Wells were coated with Speed Coating solution (500 µl per well) for 15 min at room temperature. Cells were seeded at 20,000 cells per well in complete human smooth muscle cell growth medium. At 70% confluency, cells were starved for 4 hours in OptiMEM + 2% FBS. Then, 10 nM of the respective siRNA (Silencer Select ThermoFisher) or scrambled control siRNA (Silencer Select negative control no. 2, 4390847 Thermo Fisher) was added using RNAiMAX (13778-150, Thermo Fisher). The exact protocols for all siRNA experiments are displayed below:

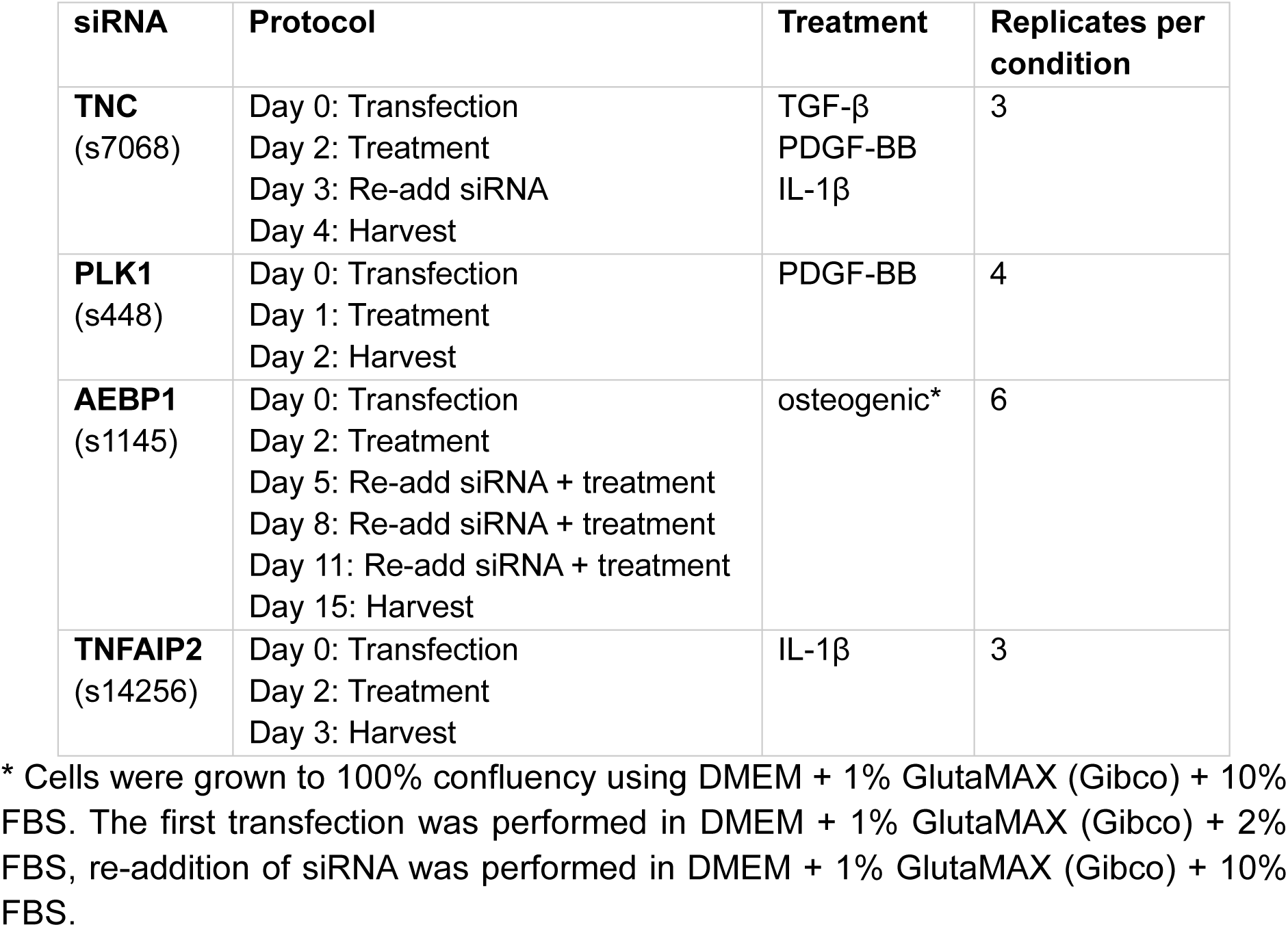

| siRNA | Protocol | Treatment | Replicates per condition |
| --- | --- | --- | --- |
| **TNC**<br>(s7068) | Day 0: Transfection<br>Day 2: Treatment<br>Day 3: Re-add siRNA<br>Day 4: Harvest | TGF-β<br>PDGF-BB<br>IL-1β | 3 |
| **PLK1**<br>(s448) | Day 0: Transfection<br>Day 1: Treatment<br>Day 2: Harvest | PDGF-BB | 4 |
| **AEBP1**<br>(s1145) | Day 0: Transfection<br>Day 2: Treatment<br>Day 5: Re-add siRNA + treatment<br>Day 8: Re-add siRNA + treatment<br>Day 11: Re-add siRNA + treatment<br>Day 15: Harvest | osteogenic* | 6 |
| **TNFAIP2**<br>(s14256) | Day 0: Transfection<br>Day 2: Treatment<br>Day 3: Harvest | IL-1β | 3 |

\* Cells were grown to 100% confluency using DMEM + 1% GlutaMAX (Gibco) + 10% FBS. The first transfection was performed in DMEM + 1% GlutaMAX (Gibco) + 2% FBS, re-addition of siRNA was performed in DMEM + 1% GlutaMAX (Gibco) + 10% FBS.

*Sample preparation* – Culture medium was collected for secretome analysis while cell pellets were collected for cellular proteomics analysis. Cell pellets were thawed prior to adding lysis buffer (60 mM TEAB, 10% acetonitrile, 10 mM DTT, 40 mM CAA). 200 µl were used for the initial VSMC treatment samples and 100 µl for siRNA-mediated knockdown samples. After heating for 30 min at 76 °C and 1000 rpm, sonication using the Bioruptor system was performed (15 cycles, 30 seconds on/off, high power). Samples were heated again for 15 min at 76 °C and 1000 rpm. After centrifugation for 5 min at 2,000 g, supernatants were transferred to new tubes. Trypsin (Sigma) and LysC (Wako) were added in a ratio of 1:50 and samples were incubated overnight (16 h) at 37 °C and 1200 rpm. Digestion was stopped by adding FA to 1% final concentration. Samples were stored at –20 °C until acquisition. Prior to LC-MS/MS acquisition, 200 ng for each sample were loaded onto Evotips. Culture medium samples were concentrated using an Amicon Ultra centrifugal filter device with a 3 kDa cut-off. Then, samples were treated similarly to the cell pellet samples. Due to the presence of phenol red in the medium of the osteogenic samples, acetone precipitation was performed prior to addition of lysis buffer. Samples were stored at –20 °C until acquisition. Prior to LC-MS/MS acquisition, 200 ng for each sample were loaded onto Evotips.

*Mass spectrometry acquisition* – Peptide samples for the primary VSMC atlas were separated by the Evosep One LC system using the 60 SPD gradient connected to an Orbitrap Astral (Thermo). An Aurora Rapid column of 8 cm length, 150 µm internal diameter, packed with 1.7 µm C18 beads (IonOpticks AUR4-8075C18-XT) was used (50 °C). Electrospray voltage was set to 1,900 V and a FAIMS Pro interface with compensation voltage of –40 V was used. Data-independent acquisition was performed over a scan range of 380-980 m/z with 3 Th windows and 6 ms maximum injection time. For the siRNA-mediated knockdown experiments, an Evosep Eno was used and the FAIMS outer electrode temperature was adjusted to 80 °C.

### Raw data analysis

Raw files were searched using a library-free search in DIA-NN (v.1.8.1.). Human FASTA files UP000005640_9606 and UP000005640_9606_additional were obtained from Uniprot (downloaded in July 2024). Default settings were used except for the following adaptions: Carbamidomethylation was disabled for the DVP samples, methionine oxidation and N-terminal protein acetylation were set as variable modifications, allowing up to one variable modification. Match-between-runs (MBR) was also enabled. Where specified, LC-MS-specific parameters were optimized on representative runs with unrelated runs option checked.

*DVP of different cell types* – Individual replicates for each cell type were searched together. Report files of each cell type were combined and protein intensities were normalized using directLFQ^43^, allowing quantitative comparison across cell types.

*DVP of VSMC neighborhoods* – Mass accuracy was set to 13, MS1 accuracy to 2 and scan window to 5.

*Primary VSMCs* – Mass accuracy was set to 7, MS1 accuracy to 5 and scan window to 6.

### Data filtering & preparation

*DVP of different cell types* – Samples with less than 1,000 PGs were excluded. The data set was filtered for 50% valid values in at least one cell type. For PCA, missing values were imputed based on a normal distribution with a width of 0.3 and downshift of 1.8.

*DVP of VSMC neighborhoods-* Samples with less than 1,000 quantified protein groups were excluded from subsequent analysis. A two-stage filtering strategy was then implemented. First, proteins were retained within individual plaques only if they were detected in at least one-third of the analyzed neighborhoods. Following this initial filtering, the resulting 24 data matrices were combined, and a second filtering criterion was applied whereby proteins were required to be detected in a minimum of 8 out of 24 plaques to be retained for downstream analysis. Missing protein values were imputed using a Python-based adaptation of the multivariate imputation by chained equations (MICE) approach^17,44^.

*Primary VSMCs* – For the cellular proteome dataset, proteins were retained if they contained at least three valid values within a single treatment group, with the requirement that these values originated exclusively from either control or treated samples. The secretome data underwent treatment-specific filtering. Cholesterol-treated samples, which showed increased protein coverage due to the absence of FBS, were filtered separately from all other treatment conditions. Proteins were retained if they contained at least three valid values within either the control or treated condition. For all remaining treatments, the same filtering criteria applied to the cellular proteome were used. Missing values in both datasets were imputed using the methodology described above. Cellular proteome and secretome datasets were merged for joint downstream analysis. Only proteins quantified across all five phenotype conditions were retained, keeping cellular and secreted entries separate to preserve compartment-specific regulation. For proteins detected in both compartments, the entry with the largest absolute log₂ fold change across treatments was retained. Per-treatment log₂ fold changes were assembled into a single protein × treatment matrix.

*siRNA-mediated knockdown samples* – For each dataset, proteins were retained if they were detected in at least half of the samples in each condition. For PCA, missing values were imputed based on a normal distribution with a width of 0.3 and downshift of 1.8. For all other downstream analysis, missing values were not imputed.

### Statistical analysis

Data analysis and visualizations were performed using custom notebooks in Python v.3.12.11, GraphPad Prism v.10.6.0 and Perseus v.2.0.10.0. Protein intensities were log_2_ transformed for all downstream analysis. Pair-wise comparisons were performed using a two-sided t-test with permutation-based FDR of 0.05.

*Subcellular compartment analysis –* The filtered dataset was intersected with GOCC gene sets retrieved from MSigDB. Ten functional compartments were defined as unions of manually selected GOCC sub-terms. Each protein group was assigned to a single compartment; protein groups matching multiple compartments were assigned to the smallest one. Proteins not matching any of the ten compartments were grouped as “Other”. Because BLOOD_MICROPARTICLE includes abundant cytoskeletal proteins expected to be overwhelmingly tissue-rather than blood-derived, we computed a sharpened composition in which any protein present in ACTIN_CYTOSKELETON, MICROTUBULE_CYTOSKELETON, INTERMEDIATE_FILAMENT_CYTOSKELETON or POLYMERIC_CYTOSKELETAL_FIBER was excluded from the blood compartment and re-assigned to its next-smallest matching compartment. Per-compartment signal was computed as the sum of linear-scale intensities of assigned proteins, expressed as percentage of total intensity. Compartment contributions to each principal component were computed as the sum of squared protein loadings per compartment divided by total PC variance.

*Gene-set enrichment analysis (GSEA)* – GSEA was performed against the Human_GOBP_AllPathways_noPFOCR_no_GO_iea_March_01_2025_symbol.gmt collection with gene set size restricted to 15-500 and 2,000 permutations^45^. To determine median activity status of biological functional terms, the median intensity of proteins belonging to the enriched pathway was calculated. Pathways with FDR q < 0.01 were retained for network visualization with the EnrichmentMap app in Cytoscape, related pathways were grouped and annotated by AutoAnnotate v1.2 based on shared keywords^46^. Single sample GSEA (ssGSEA) scores were computed for each neighborhood-term pair using the GenePattern module against GOBP, Reactome and WikiPathways gene sets.

*Over-representation analysis (ORA)* – ORA of biological processes was performed using g:Profiler^47^ with a user-defined background (all proteins after filtering, except stated otherwise) and Benjamini-Hochberg correction with FDR = 0.05 (unless stated otherwise).

*Spatial score –* For each plaque, polygon data were parsed from XML files, and the geometric centroid of each shape was computed. The luminal center was defined as the mean centroid of all shapes in the plaque, with manual x/y offsets applied where required to account for lumen geometry and position. The surrounding tissue space was divided into 120 equal angular sectors spanning 360° total. Within each sector, every centroid’s radial position was normalized using the closest and farthest centroids in that sector, yielding a spatial score in which 0 corresponds to the luminal edge and 1 to the peripheral border. Per-niche spatial scores were summarized as the mean and median across constituent centroids.

*Spatial pathway analysis –* Neighborhood samples were stratified into five equal-width spatial bins based on their spatial score. For each pathway, score differences across the five bins were tested using the Kruskal-Wallis test. Pathways with p < 0.05 were further evaluated by Dunn’s post-hoc test to identify which spatial bins differed.

*Hierarchical clustering –* Hierarchical clustering was performed using Euclidean or Pearson (for Extended Figure 5c) distance and average as linkage method. The data was pre-clustered using k-means with 300 clusters.

*Ingenuity Pathway Analysis (IPA) –* Expression analysis was performed using log_2_ fold changes to calculate directionality (z-scores) in the analysis (Cut-offs: log_2_ fc > 0.5 and < –0.5 and p-value < 0.05). Network plots of upstream regulators were created within IPA.

*Conditional variational autoencoder (CVAE) for phenotype mapping –* To integrate the in vitro VSMC reference proteomes with the VSMC neighborhood proteomes, we used a CVAE model designed to learn a shared, low-dimensional phenotype representation while minimizing information related to the sample/domain of origin. The merged cellular proteome and secretome matrix and the filtered DVP neighborhood matrix were intersected on shared proteins (n = 4,727). For each shared protein, a molecular co-expression network was constructed independently in each dataset using Spearman correlation. Proteins with similar networks (ρ > 0.3) were retained as the input feature set for CVAE (n = 1,015).

Each cell or neighborhood proteome was represented as a normalized protein-abundance vector x ∈ ℝ^p^, where p denotes the number of measured protein features. A domain indicator c, corresponding to the origin of the profile, for example *in vitro* VSMCs or DVP neighborhood, was provided as a conditional variable to the model.

The encoder network g_θ_(x, c) mapped each input proteome and its domain label to the parameters of an approximate posterior distribution over the latent phenotype embedding, q_θ_(z ∣ x, c). Specifically, the encoder produced a mean vector μ and variance parameter σ, from which latent variables were sampled using the reparameterization trick, z = μ + σ ⊙ ɛ, with ɛ ∼ *N*(0, I). The decoder network f**_φ_**(z, c) then reconstructed the input proteome from the latent embedding and the conditional domain label, yielding x⍰. Providing the domain label to the decoder allowed domain-associated variation to be modeled explicitly, thereby encouraging the latent variable z to capture phenotype-relevant variation rather than origin-specific technical or biological differences.

CVAE was trained using a composite objective containing four terms: reconstruction error, latent-space regularization, domain-invariance regularization, and sparsity regularization. Reconstruction accuracy was enforced using mean squared error between the observed and reconstructed proteome profiles. The latent distribution was regularized against a standard normal prior using the Kullback–Leibler divergence. To reduce dependence of the latent embedding on the domain of origin, we included an adversarial discriminator *k_ω_*(*z*) that attempted to predict the origin label from the latent variable *z*. The discriminator was trained to classify the origin of each profile from its latent embedding using a cross-entropy loss, whereas the encoder was trained adversarially to prevent accurate origin prediction. This adversarial objective served as a proxy for minimizing the mutual information between *z* and the domain label, thereby encouraging the latent embedding to retain proteomic variation relevant for reconstruction and phenotype mapping while reducing domain-predictive information distinguishing the in vitro VSMC reference proteomes from the in vivo VSMC neighborhood proteomes. Finally, an *L*_1_ penalty was applied to the latent representation to promote sparse phenotype encoding.

The overall CVAE objective was formulated as a minimax problem in which the encoder and decoder minimized reconstruction, KL, and sparsity losses while adversarially maximizing the domain-classification loss, and the discriminator minimized the domain-classification loss.

*Phenotype prediction of DVP neighborhoods* – The latent embedding of each DVP neighborhood was used for prediction of its similarity to the five *in vitro* VSMC reference phenotypes. Briefly, a random forest classifier implemented in scikit-learn was trained using the latent representations of the *in vitro* VSMCs as input features and their corresponding reference phenotype annotations as class labels. Hyperparameter optimization was performed using the Bayesian optimization framework Hyperopt, with searches performed over the number of trees (*n*_*estimators*), maximum tree depth (*max*_*dept*ℎ), and minimum number of samples required at each terminal leaf node (*min*_*samples*_*leaf*). After optimization, the best-performing random forest model was refit using the full *in vitro* reference dataset and applied to the latent embeddings of the DVP neighborhoods. For each DVP neighborhood, the classifier returned a probability score for each of the five reference phenotypes. These probabilities were used as phenotype-association scores, allowing each neighborhood to be represented as a weighted combination of reference VSMC phenotypic states rather than being assigned exclusively to a single phenotype.

*Comparison of phenotype distributions* – VSMC phenotype compositions were compared between stable and unstable plaques using per-patient phenotype proportions (the percentage of neighborhoods assigned to each phenotype within each patient), with patient as the unit of analysis. Group differences were assessed by a two-sided permutation test on the difference in mean per-patient proportions (10,000 permutations of the stability labels). Comparisons were not corrected for multiple testing and are reported as hypothesis-generating.

## Data availability

The mass spectrometry proteomics data, imaging data, code used and extended data tables will be available upon publication.

## Acknowledgements

We acknowledge all patients for their consent and the Munich Vascular Biobank (MVB, TUM, Munich). We thank all members of the Department of Proteomics and Signal Transduction (Max Planck Institute of Biochemistry, MPI-B, Munich) and the surgeons and research nurses of the Department for Vascular and Endovascular Surgery (TUM, Munich).

## Author contributions

EK, AS, LM and MM conceptualized the project. NS, LM and DB designed the plaque cohort. EK, AS, NS, AH and AM conducted the DVP experiments. EK, TA, VP, JP, NG and SS carried out the *in vitro* experiments. EK and TH performed mass spectrometry measurements. EK and AS performed statistical and bioinformatic data analysis. AS developed CVAE, TB and SH developed the spatial scoring system. EK, LM and MM wrote the manuscript. All authors contributed feedback on the project, data interpretation and the manuscript.

---

**Extended Figure 1:**
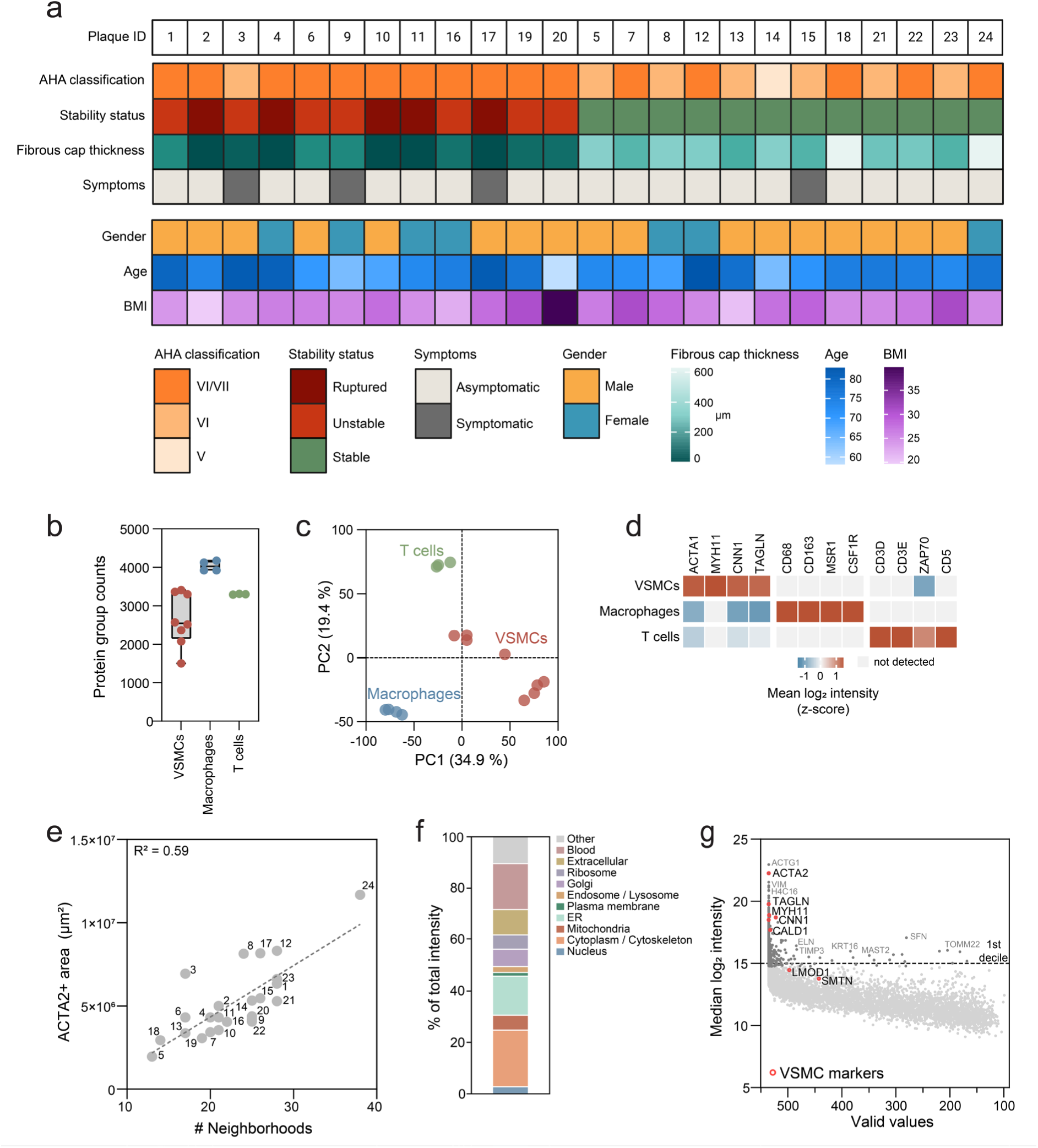
Cohort overview and DVP workflow validation. **a** Clinical covariates of patients of the 24 plaques (12 stable, 12 unstable) analyzed in this study. **b** Protein group counts of cell type benchmark after applying sample-level quality filtering (VSMCs n = 8, Macrophages n = 4, T cells n = 3). 400 shapes were dissected per sample. **c** PCA of cell type proteomes from b. **d** Heatmap of canonical cell type markers across the three cell types. Color indicates z-scored median log_2_ intensity. **e** Total ACTA2+ area versus number of dissected neighborhoods per plaque. Plaque IDs are labeled. Dashed line indicates simple linear regression. **f** Subcellular compartment distribution of the filtered proteome, aggregated as the percentage of total summed intensity across all neighborhoods. **g** Median log_2_ intensity vs number of valid values per protein group after sample– and protein-level filtering (535 neighborhoods, 5,287 protein groups). Hallmark VSMC markers are highlighted in red. The dashed line marks the 1^st^ decile of median log_2_ intensity.

**Extended Figure 2:**
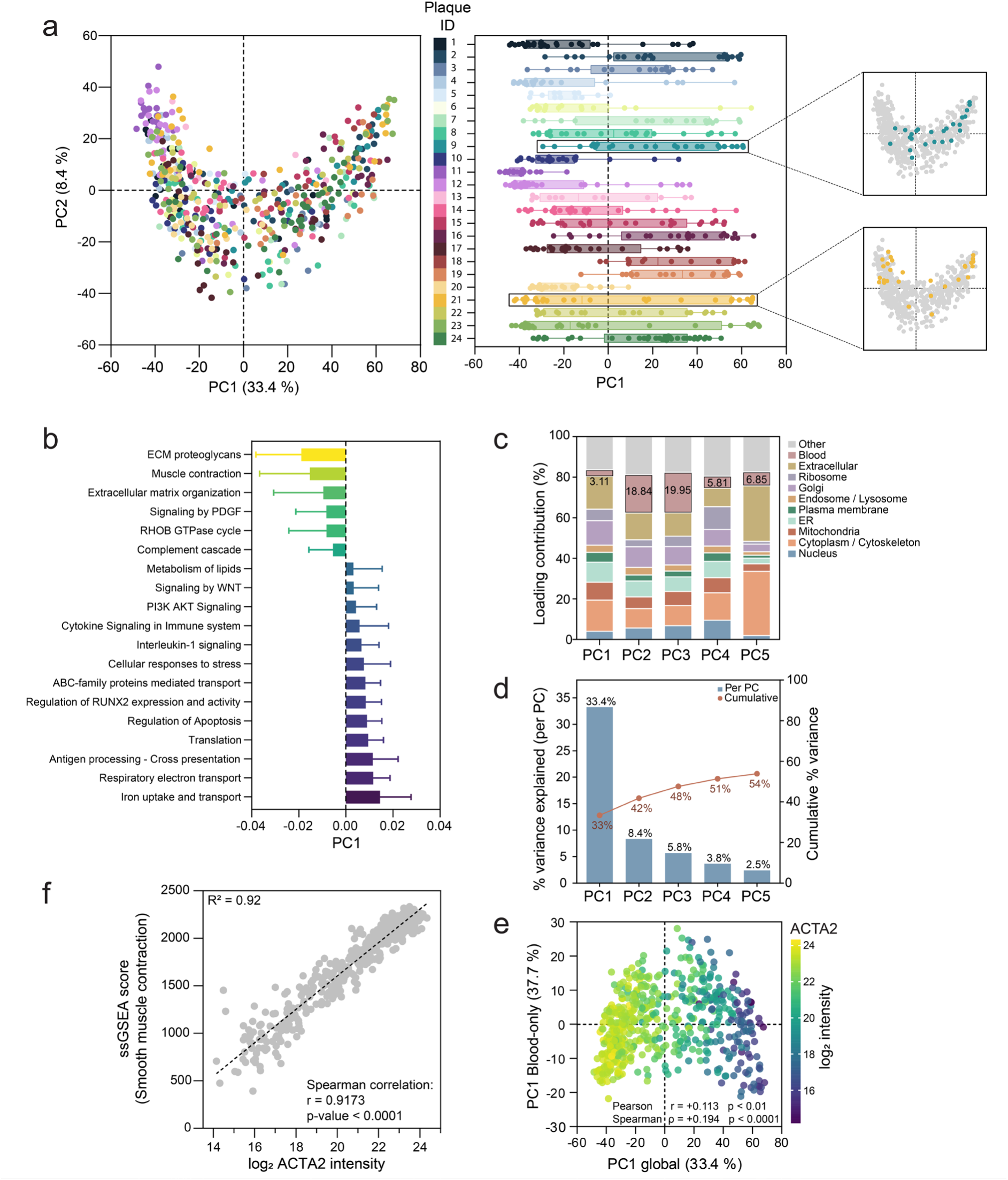
PC1 captures VSMC contractile-to-dedifferentiated continuum. **a** PCA of VSMC neighborhood proteomes (n = 535), colored by plaque ID. Left: PC1 vs PC2. Right: PC1 distributions per plaque. **b** Enrichment analysis using PC1 loadings as the ranking metric. Selected Reactome terms are shown and colored by median PC1 value. **c** Subcellular compartment contributions to principal components 1-5. **d** Explained variance per principal component 1-5 and cumulative variance. **e** PC1 computed on blood-only proteins plotted against PC1 from the global PCA, colored by ACTA2 log_2_ intensity. Pearson and Spearman correlations are reported. **f** ssGSEA score for “Smooth muscle contraction” versus ACTA2 log_2_ intensity. Dashed line indicates simple linear regression. Spearman correlation is reported.

**Extended Figure 3:**
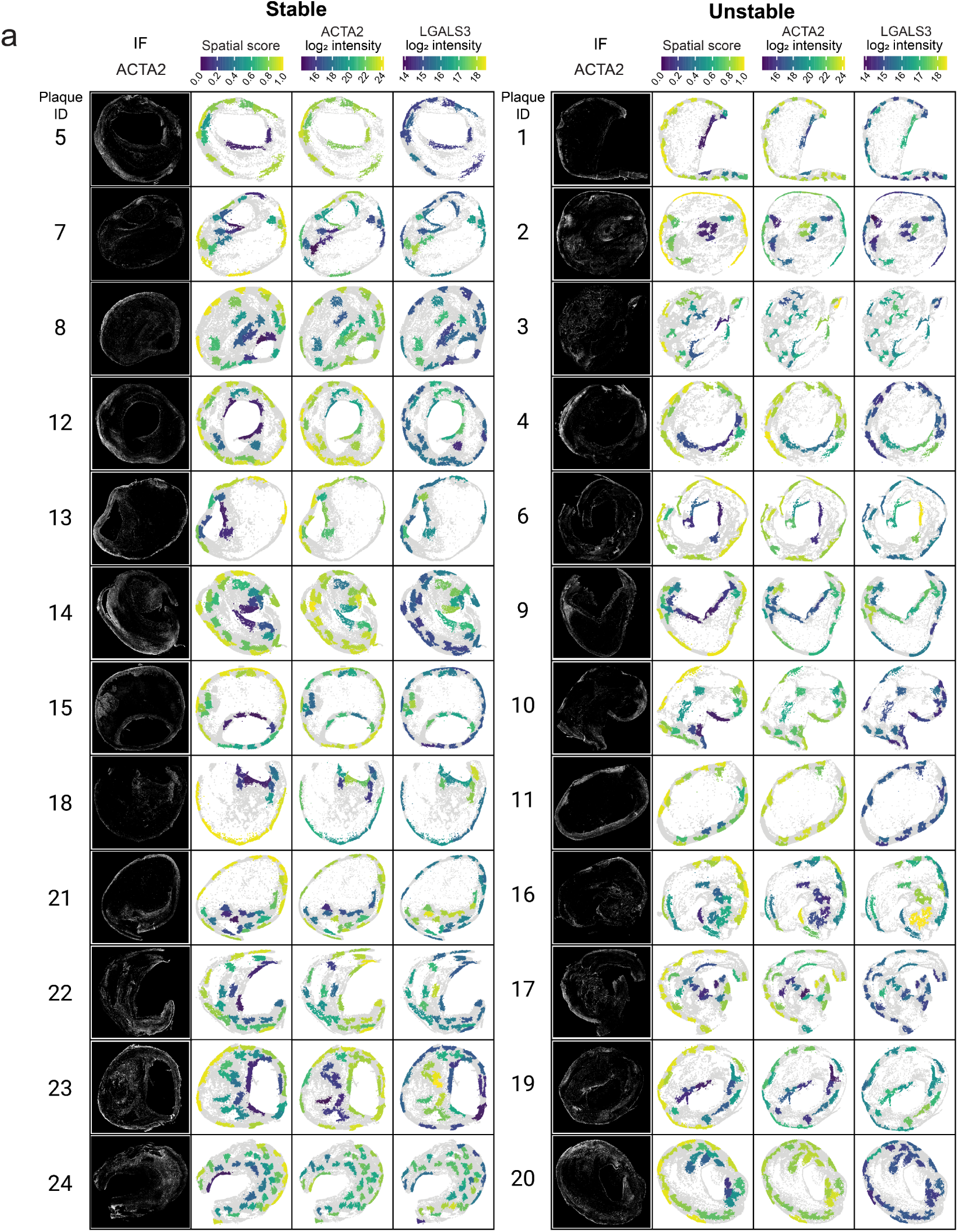
Spatial overlays for all 24 plaques. **a** ACTA2 immunofluorescence images and ACTA2+ shapes colored by spatial score, ACTA2 or LGALS3 log_2_ intensity, shown for all 24 plaques.

**Extended Figure 4:**
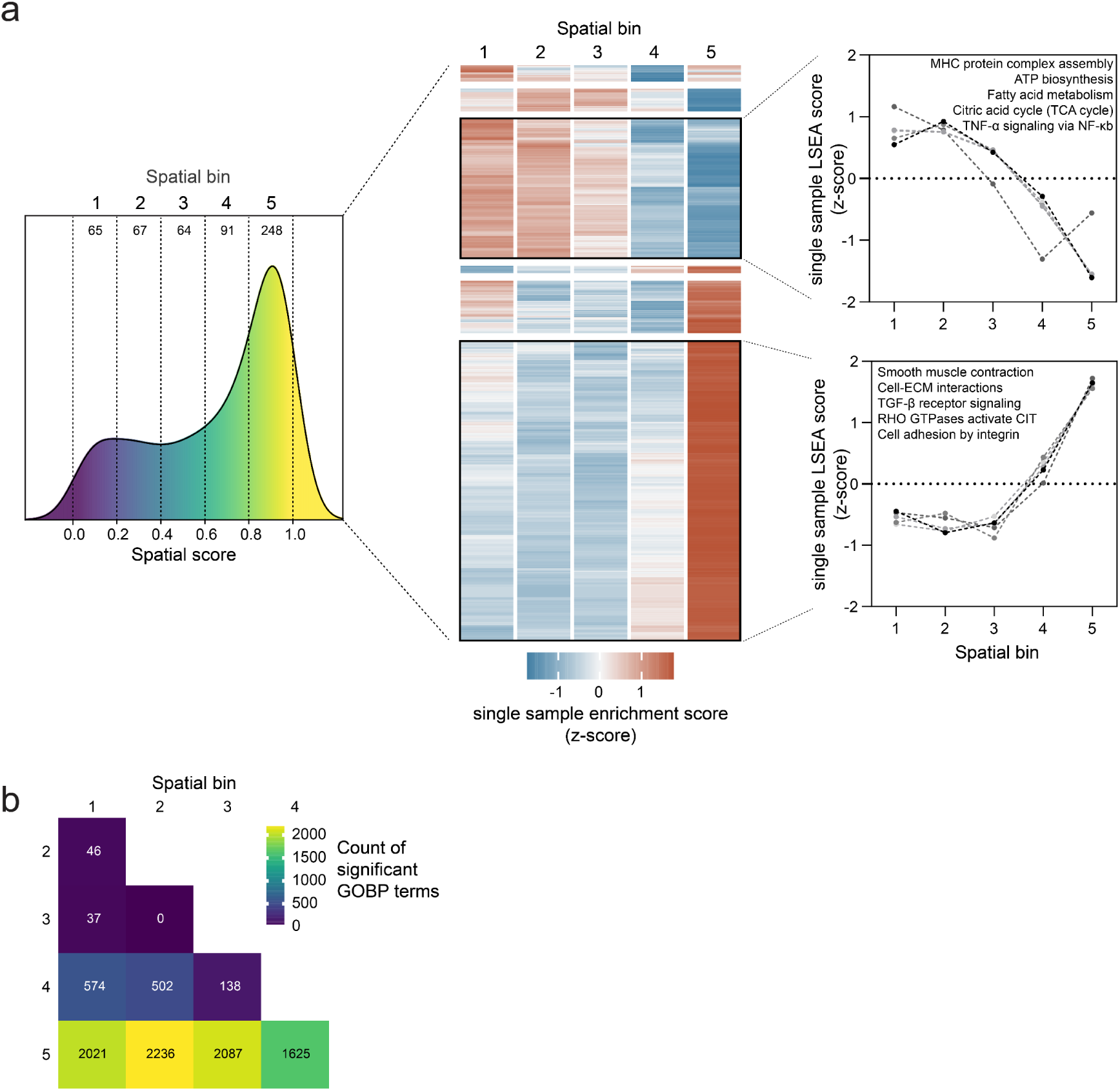
Pathway analysis across spatial score bins. **a** Neighborhoods were stratified into five spatial bins, n per spatial bin is indicated (left). Single-neighborhood enrichment scores were computed against GOBP, and pathways differing significantly across bins (Kruskal-Wallis, p < 0.05) were identified. Heatmap (center) displays all significantly differing pathways (row-based hierarchically clustered), color indicates z-scored single-sample enrichment score. Representative pathways with bin-wise enrichment trajectories are shown on the right. **b** Pairwise count of GOBP terms differing significantly between spatial bins (Dunn’s post-hoc test, p < 0.05).

**Extended Figure 5:**
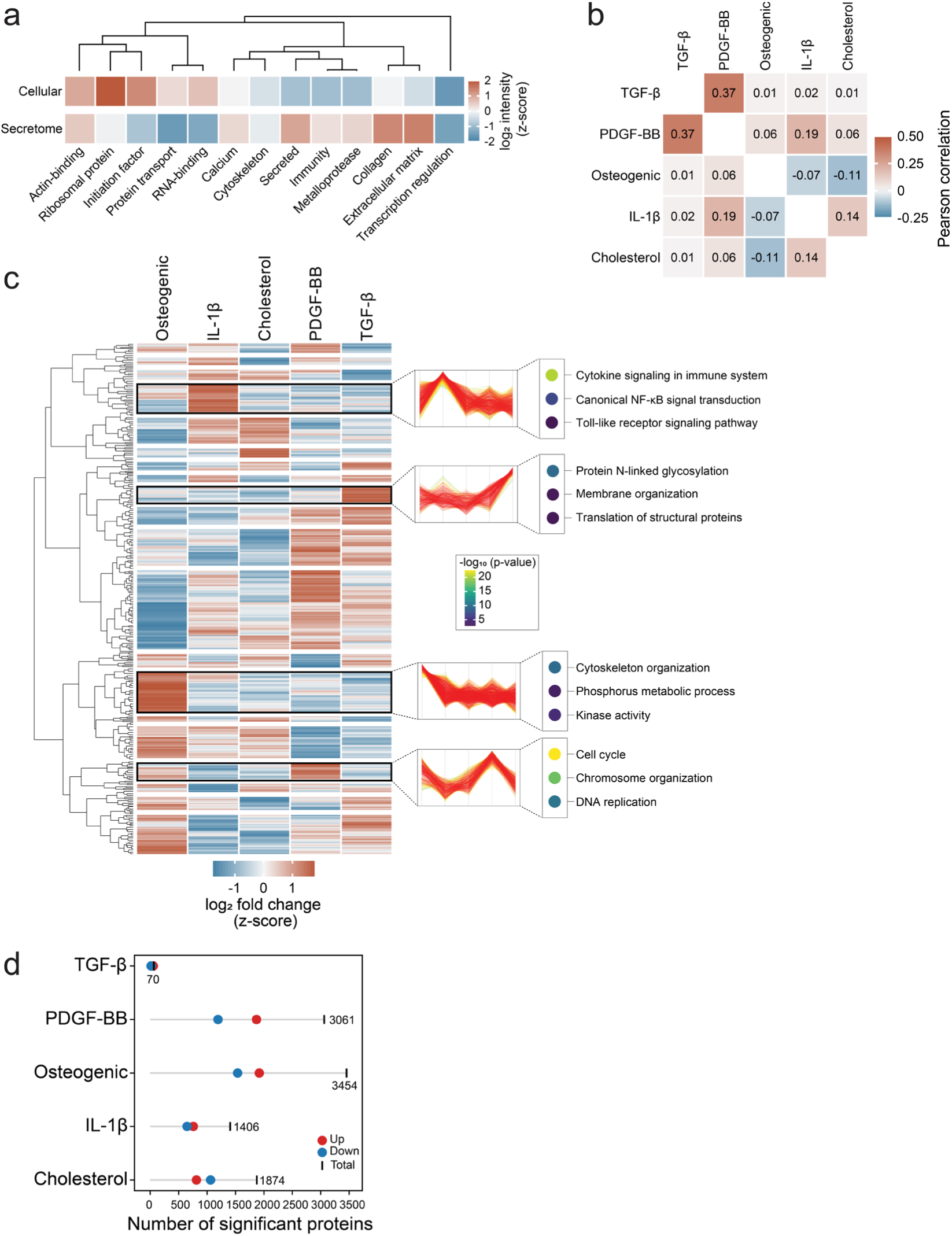
Comparison of phenotype-induced proteome signatures. **a** Mean log_2_ intensity of selected functional categories (UniProt keywords) in the cellular proteome and secretome. Columns are hierarchically clustered, color indicates z-scored mean log_2_ intensity per category. **b** Pearson correlation matrix of treatment-vs-control log_2_ fold changes across the five phenotypes, cellular proteome is shown. **c** Hierarchical clustering of treatment-vs-control log_2_ fold changes across all five phenotypes, cellular proteome is shown. Over-representation analysis of representative protein clusters and their treatment-wise trajectories are shown on the right, color indicates –log_10_ p-value. **d** Number of significantly differentially expressed proteins per treatment (up– and down-regulated, same testing as Figure 4f).

**Extended Figure 6:**
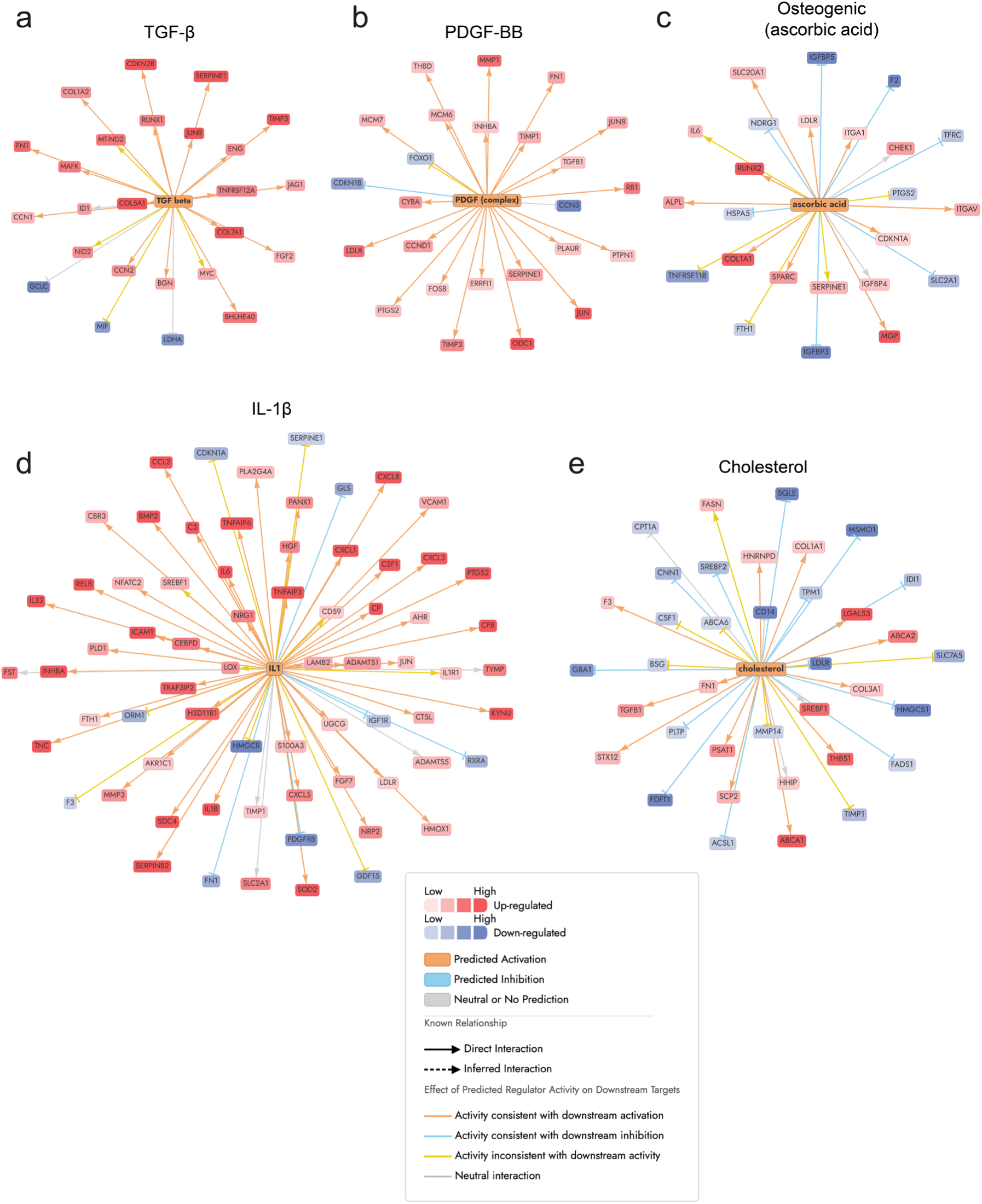
Upstream regulator analysis of phenotypic treatments. Network plots of selected upstream regulators identified by Ingenuity Pathway Analysis (IPA) for each phenotypic treatment, activation z-score and significance are indicated: **a** TGF-β (3.38, p-value = 2.80E-10). **b** PDGF complex (4.48, p-value = 4.08E-10). **c** Ascorbic acid (2.50, p-value = 1.5E-06). **d** IL1 family (5.47, p-value = 5.46E-31). **e** Cholesterol (3.60, p-value = 5.25E-08).

**Extended Figure 7:**
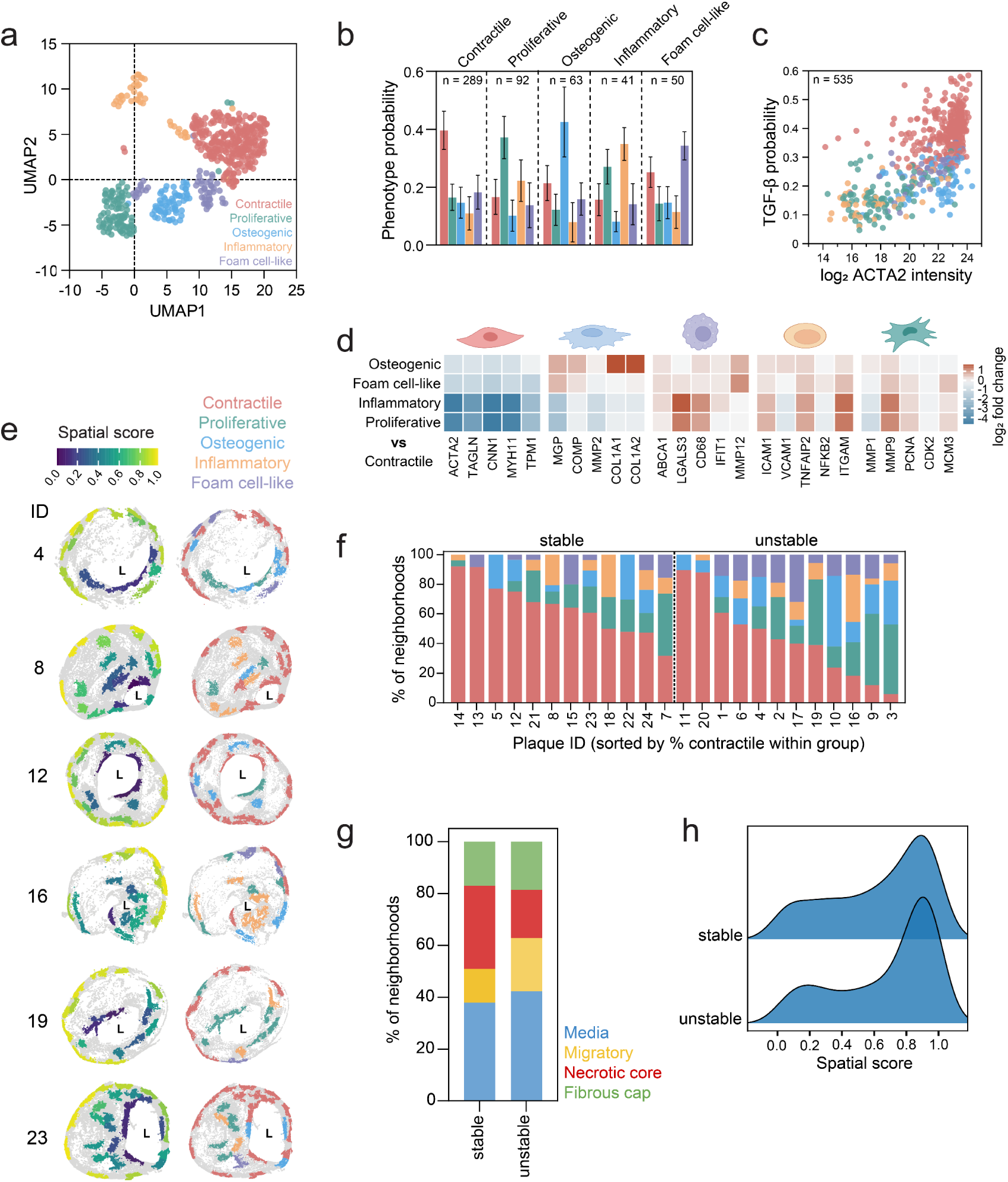
Exploration of CVAE phenotype assignments. Each neighborhood is represented by the phenotype with the highest probability. **a** UMAP (number of neighbors = 15, minimum distance = 1) of CVAE latent embeddings of VSMC neighborhood proteomes (n = 535), neighborhoods are colored by assigned phenotype. **b** Mean probability composition per assigned phenotype (mean ± SD). **c** Contractile (TGF-β) probability versus log_2_ ACTA2 intensity, neighborhoods are colored by assigned phenotype. **d** Heatmap of phenotypic markers, color indicates log_2_ fold change of each assigned dedifferentiated state versus the contractile state. **e** ACTA2+ shapes colored by spatial score (left) and phenotype identity (right), shown for plaques 4, 8, 12, 16, 19 and 23. L = plaque lumen. **f** Phenotype proportions per plaque reflecting neighborhood-level counts, sorted by descending percentage of contractile neighborhoods within each group. **g** Subregion composition of stable and unstable plaques. **h** Density distribution of spatial scores in stable and unstable plaques.

**Extended Figure 8:**
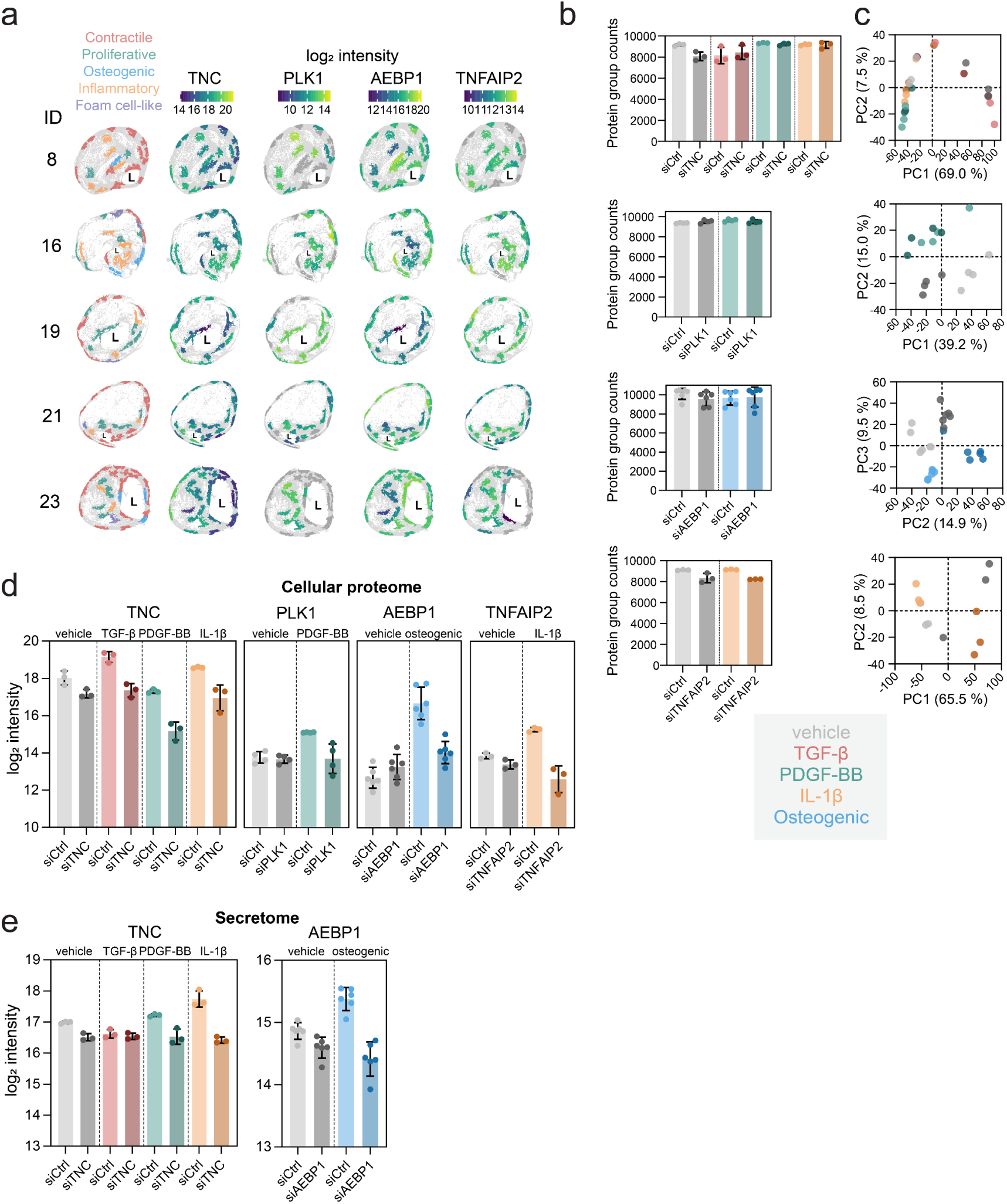
Quality control of perturbation experiments. Each neighborhood is represented by the phenotype with the highest probability. **a** ACTA2+ shapes colored by phenotype identity (left) or by log_2_ intensity of TNC, PLK1, AEBP1 and TNFAIP2 (right). Shown for plaques 8, 16, 19, 21 and 23. L = plaque lumen. **b** Protein group counts per condition (cellular proteome). **c** PCA of cellular proteomes, colored by siRNA and treatment condition. **d** Log_2_ intensity of TNC, PLK1, AEBP1 and TNFAIP2 in respective conditions in cellular proteome. **e** Log_2_ intensity of TNC and AEBP1 in respective conditions in secretome.

## References

1. Stark, B. A. et al. Global, Regional, and National Burden of Cardiovascular Diseases and Risk Factors in 204 Countries and Territories, 1990-2023. JACC 86, 2167–2243 (2025).

2. Libby, P. The changing landscape of atherosclerosis. Nature 592, 524–533 (2021).

3. Kong, P. et al. Inflammation and atherosclerosis: signaling pathways and therapeutic intervention. Signal Transduct. Target. Ther. 7, 131 (2022).

4. Traeuble, K. et al. Integrated single-cell atlas of human atherosclerotic plaques. Preprint at 10.1101/2024.09.11.612431 (2024).

5. Bennett, M. R., Sinha, S. & Owens, G. K. Vascular Smooth Muscle Cells in Atherosclerosis. Circ. Res. 118, 692–702 (2016).

6. Lin, A., Miano, J. M., Fisher, E. A. & Misra, A. Chronic inflammation and vascular cell plasticity in atherosclerosis. *Nat*. Cardiovasc. Res. 3, 1408–1423 (2024).

7. Grootaert, M. O. J. & Bennett, M. R. Vascular smooth muscle cells in atherosclerosis: time for a re-assessment. Cardiovasc. Res. 117, 2326–2339 (2021).

8. Yap, C., Mieremet, A., De Vries, C. J. M., Micha, D. & De Waard, V. Six Shades of Vascular Smooth Muscle Cells Illuminated by KLF4 (Krüppel-Like Factor 4). Arterioscler. Thromb. Vasc. Biol. 41, 2693–2707 (2021).

9. Sorokin, V. et al. Role of Vascular Smooth Muscle Cell Plasticity and Interactions in Vessel Wall Inflammation. Front. Immunol. 11, 599415 (2020).

10. Rosenfeld, M. E. Converting smooth muscle cells to macrophage-like cells with KLF4 in atherosclerotic plaques. Nat. Med. 21, 549–551 (2015).

11. Maegdefessel, L. & Fasolo, F. Long Non-Coding RNA Function in Smooth Muscle Cell Plasticity and Atherosclerosis. Arterioscler. Thromb. Vasc. Biol. 45, 172–185 (2025).

12. Azar, P. et al. Smooth muscle cells in atherosclerosis: essential but overlooked translational perspectives. Eur. Heart J. 46, (2025).

13. Wirka, R. C. et al. Atheroprotective roles of smooth muscle cell phenotypic modulation and the TCF21 disease gene as revealed by single-cell analysis. Nat. Med. 25, 1280–1289 (2019).

14. Pan, H. et al. Single-Cell Genomics Reveals a Novel Cell State During Smooth Muscle Cell Phenotypic Switching and Potential Therapeutic Targets for Atherosclerosis in Mouse and Human. Circulation 142, 2060–2075 (2020).

15. Pauli, J. et al. Single cell spatial transcriptomics integration deciphers the morphological heterogeneity of atherosclerotic carotid arteries. Nat. Commun. 16, 11282 (2025).

16. Goncalves, I. et al. Spatial transcriptomics reveals a key role of fibroblast-like vascular smooth muscle cells in human atherosclerotic cell crosstalk and stability. Eur. Heart J. 1–22 (2026) doi:10.1093/eurheartj/ehaf1091.

17. Sinha, A. et al. Spatially resolved proteomic signatures of atherosclerotic carotid artery disease. Preprint at 10.1101/2025.02.09.25321955 (2025).

18. Mund, A. et al. Deep Visual Proteomics defines single-cell identity and heterogeneity. Nat. Biotechnol. 40, 1231–1240 (2022).

19. Redgrave, J. N., Gallagher, P., Lovett, J. K. & Rothwell, P. M. Critical Cap Thickness and Rupture in Symptomatic Carotid Plaques: The Oxford Plaque Study. Stroke 39, 1722–1729 (2008).

20. Stary, H. C. et al. A Definition of Advanced Types of Atherosclerotic Lesions and a Histological Classification of Atherosclerosis: A Report From the Committee on Vascular Lesions of the Council on Arteriosclerosis, American Heart Association. Arterioscler. Thromb. Vasc. Biol. 15, 1512–1531 (1995).

21. Aebersold, R. & Mann, M. Mass-spectrometric exploration of proteome structure and function. Nature 537, 347–355 (2016).

22. Perisic Matic, L. et al. Phenotypic Modulation of Smooth Muscle Cells in Atherosclerosis Is Associated With Downregulation of LMOD1, SYNPO2, PDLIM7, PLN, and SYNM. Arterioscler. Thromb. Vasc. Biol. 36, 1947–1961 (2016).

23. Yu, Y. et al. Vascular smooth muscle cell phenotypic switching in atherosclerosis. Heliyon 10, e37727 (2024).

24. Shankman, L. S. et al. KLF4-dependent phenotypic modulation of smooth muscle cells has a key role in atherosclerotic plaque pathogenesis. Nat. Med. 21, 628– 637 (2015).

25. Bleckwehl, T. et al. Encompassing view of spatial and single-cell RNA sequencing renews the role of the microvasculature in human atherosclerosis. Nat. Cardiovasc. Res. https://doi.org/10.1038/s44161-024-00582-1 (2024) doi:10.1038/s44161-024-00582-1.

26. Virmani, R., Kolodgie, F. D., Burke, A. P., Farb, A. & Schwartz, S. M. Lessons From Sudden Coronary Death: A Comprehensive Morphological Classification Scheme for Atherosclerotic Lesions. Arterioscler. Thromb. Vasc. Biol. 20, 1262– 1275 (2000).

27. Low, E. L., Baker, A. H. & Bradshaw, A. C. TGFβ, smooth muscle cells and coronary artery disease: a review. Cell. Signal. 53, 90–101 (2019).

28. Chen, C., Yan, Y. & Liu, X. microRNA-612 is downregulated by platelet-derived growth factor-BB treatment and has inhibitory effects on vascular smooth muscle cell proliferation and migration via directly targeting AKT2. Exp. Ther. Med. https://doi.org/10.3892/etm.2017.5428 (2017) doi:10.3892/etm.2017.5428.

29. Pustlauk, W. et al. Induced osteogenic differentiation of human smooth muscle cells as a model of vascular calcification. Sci. Rep. 10, 5951 (2020).

30. Eun, S. Y., Ko, Y. S., Park, S. W., Chang, K. C. & Kim, H. J. IL-1β enhances vascular smooth muscle cell proliferation and migration via P2Y2 receptor-mediated RAGE expression and HMGB1 release. Vascul. Pharmacol. 72, 108– 117 (2015).

31. Bogan, B. J. et al. The Role of Fatty Acid Synthase in the Vascular Smooth Muscle Cell to Foam Cell Transition. Cells 13, 658 (2024).

32. Guan, X. et al. Stress, Vascular Smooth Muscle Cell Phenotype and Atherosclerosis: Novel Insight into Smooth Muscle Cell Phenotypic Transition in Atherosclerosis. Curr. Atheroscler. Rep. 26, 411–425 (2024).

33. Imanaka-Yoshida, K., Yoshida, T. & Miyagawa-Tomita, S. Tenascin-C in Development and Disease of Blood Vessels. Anat. Rec. 297, 1747–1757 (2014).

34. Sur, S., Swier, V. J., Radwan, M. M. & Agrawal, D. K. Increased Expression of Phosphorylated Polo-Like Kinase 1 and Histone in Bypass Vein Graft and Coronary Arteries following Angioplasty. PLOS ONE 11, e0147937 (2016).

35. Layne, M. D. et al. Characterization of the Mouse Aortic Carboxypeptidase-Like Protein Promoter Reveals Activity in Differentiated and Dedifferentiated Vascular Smooth Muscle Cells. Circ. Res. 90, 728–736 (2002).

36. Jin, G. et al. Tnfaip2 promotes atherogenesis by enhancing oxidative stress induced inflammation. Mol. Immunol. 151, 41–51 (2022).

37. He, J. et al. Navigating the landscape: Prospects and hurdles in targeting vascular smooth muscle cells for atherosclerosis diagnosis and therapy. J. Controlled Release 366, 261–281 (2024).

38. Paloschi, V. et al. Utilization of an Artery-on-a-Chip to Unravel Novel Regulators and Therapeutic Targets in Vascular Diseases. Adv. Healthc. Mater. 13, 2302907 (2024).

39. Parma, L. et al. CXCL12 Derived From ACKR1^+^ Intraplaque Neovessels Mediates CD8^+^ T Cell Recruitment in Human Atherosclerosis. Circulation 151, 581–584 (2025).

40. Bentzon, J. F. & Falk, E. Atherosclerotic lesions in mouse and man: is it the same disease?: Curr. Opin. Lipidol. 21, 434–440 (2010).

41. Worssam, M. D. & Jørgensen, H. F. Mechanisms of vascular smooth muscle cell investment and phenotypic diversification in vascular diseases. Biochem. Soc. Trans. 49, 2101–2111 (2021).

42. Bankhead, P. et al. QuPath: Open source software for digital pathology image analysis. Sci. Rep. 7, 16878 (2017).

43. Ammar, C., Schessner, J. P., Willems, S., Michaelis, A. C. & Mann, M. Accurate Label-Free Quantification by directLFQ to Compare Unlimited Numbers of Proteomes. Mol. Cell. Proteomics 22, 100581 (2023).

44. Buuren, S. V. & Groothuis-Oudshoorn, K. mice: Multivariate Imputation by Chained Equations in R. J. Stat. Softw. 45, (2011).

45. Subramanian, A. et al. Gene set enrichment analysis: A knowledge-based approach for interpreting genome-wide expression profiles. Proc. Natl. Acad. Sci. 102, 15545–15550 (2005).

46. Reimand, J. et al. Pathway enrichment analysis and visualization of omics data using g:Profiler, GSEA, Cytoscape and EnrichmentMap. Nat. Protoc. 14, 482– 517 (2019).

47. Reimand, J., Kull, M., Peterson, H., Hansen, J. & Vilo, J. g:Profiler—a web-based toolset for functional profiling of gene lists from large-scale experiments. Nucleic Acids Res. 35, W193–W200 (2007).

48. Perez-Riverol, Y. et al. The PRIDE database at 20 years: 2025 update. Nucleic Acids Res. 53, D543–D553 (2025).

49. Sarkans, U. et al. The BioStudies database—one stop shop for all data supporting a life sciences study. Nucleic Acids Res. 46, D1266–D1270 (2018).

